# Antigen stimulation drives clonal expansion of latent CD4^+^ T cells using a full-length HIV latency reporter

**DOI:** 10.1101/2025.09.04.672206

**Authors:** Nnamdi Ikeogu, Oluwaseun Ajibola, Riley Greenslade, Wan Koh, Amélie Pagliuzza, Rémi Fromentin, Michelle Perner, Roshan Parvarchian, Xinyun Liu, Paul Lopez, Catherine Card, Paul McLaren, Nicolas Chomont, Mario Ostrowski, Thomas Murooka

**Author notes:** Contributed equally to this study (Oluwaseun Ajibola, Riley Greenslade). **Address correspondence to:** Thomas T. Murooka, Department of Immunology, Rady Faculty of Health Sciences, University of Manitoba, Canada. **Authorship:**. Contributions: NI, OA, RG and TTM conceived and designed the experiments; NI, OA, RG, AP, RP, XL, PL and CC performed the experiments; WHK and MO provided CD4^+^ T cell clones; MP and PM performed TCRb sequencing and analysis; NI, OA, RG, AA, NC and TTM analyzed the data; NI, OA and TTM wrote the manuscript. **Competing interest statement:** The authors declare no competing interests.

## Abstract

HIV persistence despite years of ART suppression poses a major barrier to cure. Using a full-length latency reporter to generate HIV-infected, transcriptionally silent CD4^+^ T cells *in vitro*, we show that cognate DC:T cell interactions drive clonal expansion of latent T cells in an antigen dependent manner and that a pro-survival state within proliferating cells is reinforced through IL-7 signaling. Interestingly, we describe a dominant role for CD28 co-stimulation in regulating robust latent T cell proliferation which was partially reversed by PD-1 blockade. Our studies show that a gradual reduction in antigenic stimulation was sufficient to induce proliferative responses without measurable proviral reactivation. Thus, the magnitude of TCR/co-stimulatory signals during cognate APC:T cell interactions are key regulators of the underlying proliferative and survival programs maintaining the latent reservoir under ART suppression.

**Significance Statement:** Viral replication can be effectively suppressed by antiretroviral therapy (ART) but is not curative due to persistence of latent virus in a stable reservoir in resting CD4^+^ T cells. We show that antigen recognition through cell-cell interactions is an important driver of latent T cell proliferation, and that modulating TCR stimulatory signaling independently regulates proliferative, survival and proviral reactivation potential in infected T cells. Our observations show that latent T cells retain their ability to engage other immune cells to support their long-term survival under ART suppression, similar to uninfected T cells. These characteristics of latent T cell pools represent an additional hurdle to eradicating the reservoir.

## Introduction

Successful antiretroviral therapy (ART) can efficiently suppress Human Immunodeficiency Virus-1 (HIV-1; referred to as HIV) replication to undetectable levels but rare subsets of infected memory CD4^+^ T cells continue to persist, complicating viral eradication efforts (1–4). Establishing viral latency in memory CD4^+^ T cells offers several advantages. First, HIV-infected T cells retain their motility, albeit at reduced speeds, providing a means for the virus to disseminate widely across distant organ systems while minimizing clearance by the immune system (5–7). Cell migration and initiation of cell-cell contacts within tissues are well-established conduits for efficient local HIV spread (8–12). Second, memory T cells are long-lived and continually receive survival/proliferative signals during their physiological transit through lymphoid and non-lymphoid organs (13–15). Long-term maintenance and function of the memory T cell compartment is reliant on at least two important signals: (i) TCR signaling initiated through cell-cell contacts with antigen presenting cells (16, 17) and (ii) cytokines such as IL-7 and IL-15 that upregulate anti-apoptotic pathways and induce homeostatic proliferation to replenish the memory T cell pool (18–21). Thus, establishment of post-integration latency in memory T cells may allow access to survival cues by restoring physiological recirculation across secondary lymphoid organs (13, 22, 23).

TCR engagement with the peptide-MHC complex in response to infection results in T cell activation and clonal expansion. Emerging data show that latently infected T cells carrying replication-competent provirus also persist through clonal expansion *in vivo*. Mendoza and colleagues showed that CD4^+^ T cells containing defective or intact provirus were found in T cells responsive to HIV Gag, Cytomegalovirus (CMV), Epstein-Barr virus and Influenza virus-derived antigens (24). Similarly, Simonetti *et al* isolated CMV- and Gag-responding CD4^+^ T cells from ART-treated individuals and observed large clones harboring replication-competent provirus that were not explained by homeostatic proliferation or specific integration site effects (25). Moreover, recognition of self-antigens can also induce HIV production in an MHC-II dependent manner, indicating that both self and foreign-derived peptides can activate infected T cells (26). Recent findings suggest that the reservoir may begin to slowly increase after the first 7 years of ART suppression (27). These studies indicate that HIV persistence is largely driven by infected cell proliferation, and that repeated exposure to antigens may drive clonal expansion over time (15, 28–35). HIV DNA^+^ memory CD4^+^ T cells also display transcriptional signatures associated with inhibition of death receptor/necroptosis signaling and anti-proliferative Gα12/13 pathways, indicative of a strong proliferative program under ART (36–39). T cell clones harboring replication-competent provirus were found to be mostly transcriptionally silent at any given time, indicating that proliferation can occur in the absence of overt viral replication in ART-suppressed individuals (40, 41). However, it remains unclear how antigen specific TCR stimulation regulates seemingly opposing biological processes of infected T cell clonal expansion while avoiding reversal of latency that might lead to either cytotoxicity or clearance by the immune system. Here, we test the hypothesis that infected T cells undergo proliferation if the driving stimulus is not strong enough to upregulate HIV gene transcription. We speculate that, over the course of their transit through peripheral tissues and blood, latent T cells continually integrate tonic and antigenic signals to control a range of cellular responses, some of which support their long-term persistence.

Here, we generated and validated a new full-length, dual-fluorescent HIV reporter in primary CD4^+^ T cells to perform repeated phenotypic and proliferative analyses in *in vitro* generated latent T cells. We provide direct evidence that cognate DC:T cell interactions drive clonal expansion of latent T cell pools in the absence of proviral reactivation, and that IL-7 reinforces a pro-survival state within proliferating cells. Using primary CD4^+^ T cell clones with a fixed TCR, we show that a stepwise reduction in antigenic stimulation induced strong pro-survival and proliferative responses in the absence of measurable viral transcription. Modulation of TCR signaling strengths through PD-1 blockade favors HIV transcription and reduces the proliferative potential of CD4^+^ T cell clones. These studies argue that antigenic TCR signaling strength may regulate molecular programs that drive expansion of distinct T clones under ART suppression. Inducing a state of latency may allow infected T cells to retain their physiological capabilities to engage in cognate cell-cell interactions *in vivo* and undergo antigen-driven proliferation.

## Results

### Full length R5-tropic HIV-2A-CRIMZs reporter delineates productive and latent infection in primary CD4^+^ T cells

To visually characterize latent CD4^+^ T cells, a new full-length dual-fluorescent HIV reporter was generated that encodes a Nef-2A-Crimson gene locus under the control of the HIV LTR promoter, and ZsGreen under the control of the mammalian EF1a-HTLV promoter (called HIV Nef-2A-CRIMZs, Fig. 1A). This reporter contains all structural and accessory HIV genes and is replication competent. Constitutive expression of ZsGreen allows us to identify T cells that have stably integrated provirus, while Crimson expression reports LTR-driven transcription activity (42–44). To validate this reporter, naïve CD4^+^ T cells were isolated from donor PBMCs and activated with αCD3/CD28 dynabeads for 7 days to generate central memory-like T cells (45). Resting CD4^+^ T cells were infected with HIV Nef-2A-CRIMZs in collagen gels at an MOI of 0.1 (Fig. 1B), as described previously (46). Flow cytometry analysis at 5 days post-infection visually discriminated Crimson^+^Zsgreen^+^ cells (productive infection; orange box) from Crimson^neg^Zsgreen^+^ cells (latent infection; green box), whereas virally exposed but double-negative (DN) T cells remained Crimson^neg^Zsgreen^neg^ (Fig. 1C). Both populations were further evaluated for intracellular Gag p24 and cell surface CD4 expression, where productively infected T cells were Gag p24^+^ and displayed reduced CD4 expression, indicative of Env and Nef-mediated CD4 downregulation (Fig. 1C; Suppl. Fig. 1) (47). Notably, a higher ratio of latently infected T cells was observed with the reporter containing *Nef*, consistent with another study (48), suggesting reduced productively infected T cell frequencies over time due to Nef-mediated apoptotic death (49). In contrast, the latent T cell population remained Gag p24 negative and retained high CD4 expression. Live cell sorting at Day 3 post-infection confirmed high Gag p24 expression in productively infected T cells and no memory phenotypic differences between populations were observed, as they all expressed central memory markers (Fig. 1D). To measure proviral reactivation potential, latent T cell pools were live sorted and stimulated with anti-CD3/CD28 antibody for evaluation 48 hours later by flow cytometry. Latent T cells displayed a quiescent phenotype at baseline characterized by low CD69 and HLA-DR expression, whereas TCR stimulation significantly upregulated their expression along with Gag p24 production (Fig. 1E,F). Collectively, our HIV reporter defines latent T cells as ZsGreen^hi^CD4^hi^ and permits studies into the molecular cues that support their proliferative and survival responses under ART suppression.

**Figure 1:**
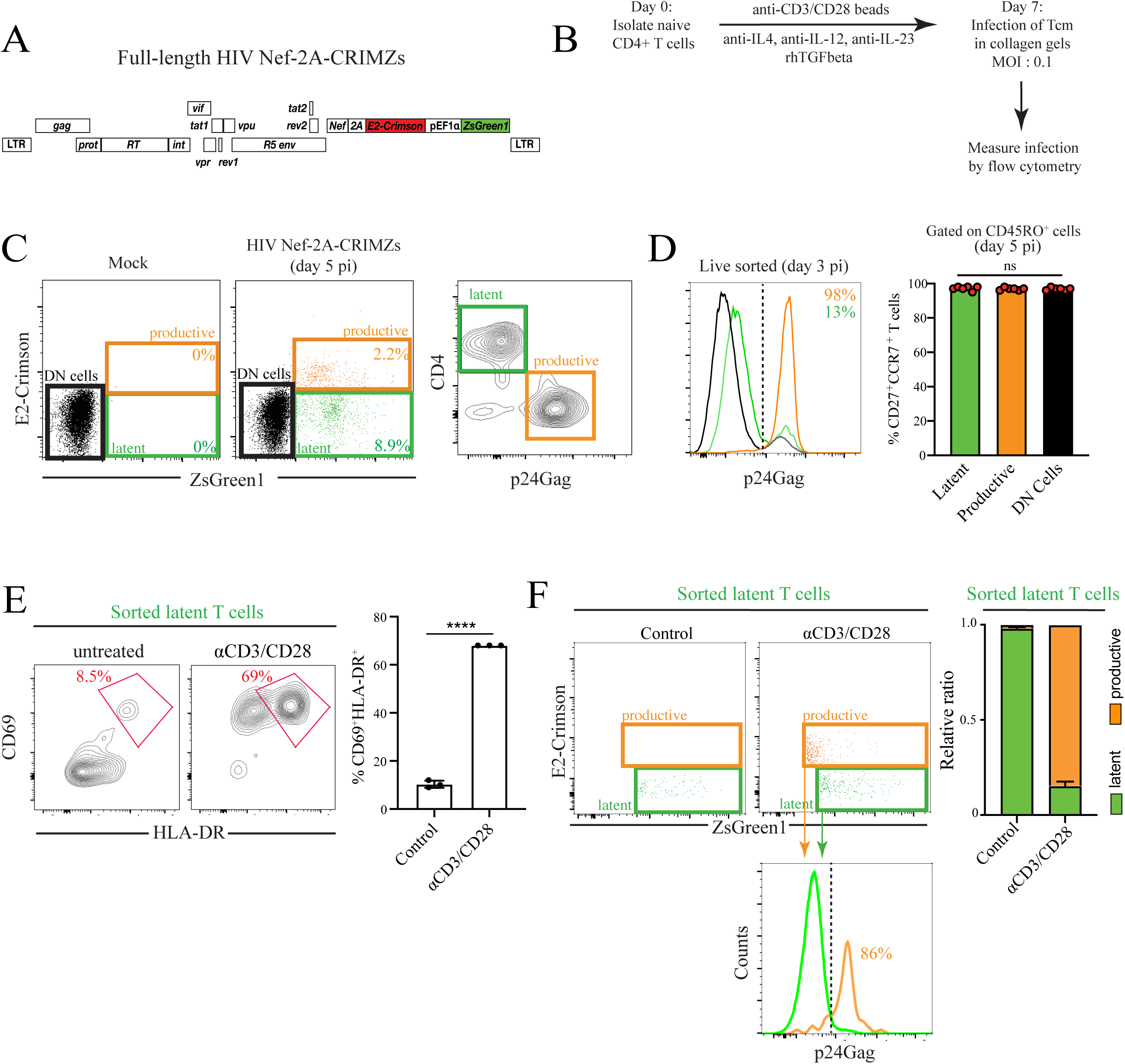
Construction and validation of the HIV Nef-2A-CRIMZs reporter in primary CD4^+^ T cells. (**A**) Schematic representation of HIV Nef-2A-CRIMZs reporter. **(B)** Naive CD4^+^ T cells from blood donors were isolated and cultured under non-polarizing condition for 7 days to generate Tcm-like cells. T cells were infected with HIV Nef-2A-CRIMZs at MOI of 0.1 in collagen gels and assessed for infection at day 5. **(C)** Distribution of productive vs latent infection using HIV Nef-2A-CRIMZs is shown using orange and green boxes, respectively, by flow cytometry. Cell surface CD4 expression across all three T cell populations is shown. Representative data from experiments performed in 6 donors. **(D)** Infected T cells (Day 3) were live cell sorted and evaluated for p24Gag expression. Central memory phenotype of latent, productive and HIV-exposed, double-negative (DN) T cells is also shown. Tukey’s multiple comparisons test. ns = not significant. **(E)** Latent T cells defined as ZsGreen^+^Crimzon^neg^CD4^hi^ were live cell sorted and stained for CD69 and HLA-DR expression before and after stimulation with aCD3/CD28 conjugated dynabeads for 24 hours. Red box and corresponding percentage depict CD69^+^HLA-DR^+^ T cells. Representative data from experiments performed in 3 donors. ns = not significant. **** p<0.0001, unpaired t-test. **(F)** Intracellular staining for Gag p24 was performed before and after stimulation. Relative ratio of productive vs latent T cell populations before and after aCD3/CD28 stimulation is shown. **** p<0.0001, unpaired t-test. Data from a representative donor shown.

### Dendritic cells and IL-7 promote proliferation and survival of latent primary CD4^+^ T cells

Tonic TCR signaling generated through transient cell-cell contacts with dendritic cells along with IL-7 are contributing factors to the long-term maintenance and function of memory T cell pools (16–19). We investigated whether similar cellular and molecular processes were involved in latent T cell persistence under ART. We first compared IL-7R expression between productively and latently infected T cells, with the latter population expressing significantly higher cell surface IL-7R (Fig. 2A). Comparatively, DN T cells expressed high IL-7R levels. Next, we performed a series of autologous DC:T cell co-culture studies in the presence or absence of IL-7 to examine their contribution to infected T cell proliferation and survival. Primary CD4^+^ T cells from healthy donors were isolated and expanded into central memory-like T cells prior to infection with HIV Nef-2A-CRIMZs, similar to Figure 1B. At day 5 post-infection, T cells were stained with a violet proliferation tracker dye (Tag-IT) and co-cultured with autologous monocyte-derived dendritic cells (MDDCs) for a further 10 days to measure recall responses in the presence of the integrase inhibitor, raltegravir to block secondary infections (Fig. 2B, see Suppl. Fig 2 for representative gating strategy) (46). We show that both DCs and IL-7 promote expansion of the latent T cell pool, with additive effects seen in the presence of both DCs and IL-7 (Fig 2C-F). IL-7 significantly upregulated anti-apoptotic bcl-2 expression (Fig. 2G). Similar proliferative and survival responses were upregulated in HIV exposed but double-negative T cells present in the same co-culture, indicating that observed responses were not unique to HIV-infected T cells (Fig. 2F,G). Importantly, we observed no HIVGag+ T cells after 4 or 10 days of co-culture, likely due to cell death and secondary infections blocked with ART. Together, we show that tonic DC:T cell interactions, together with IL-7, was sufficient to promote tonic proliferation and induce a pro-survival state in latent T cells.

**Figure 2:**
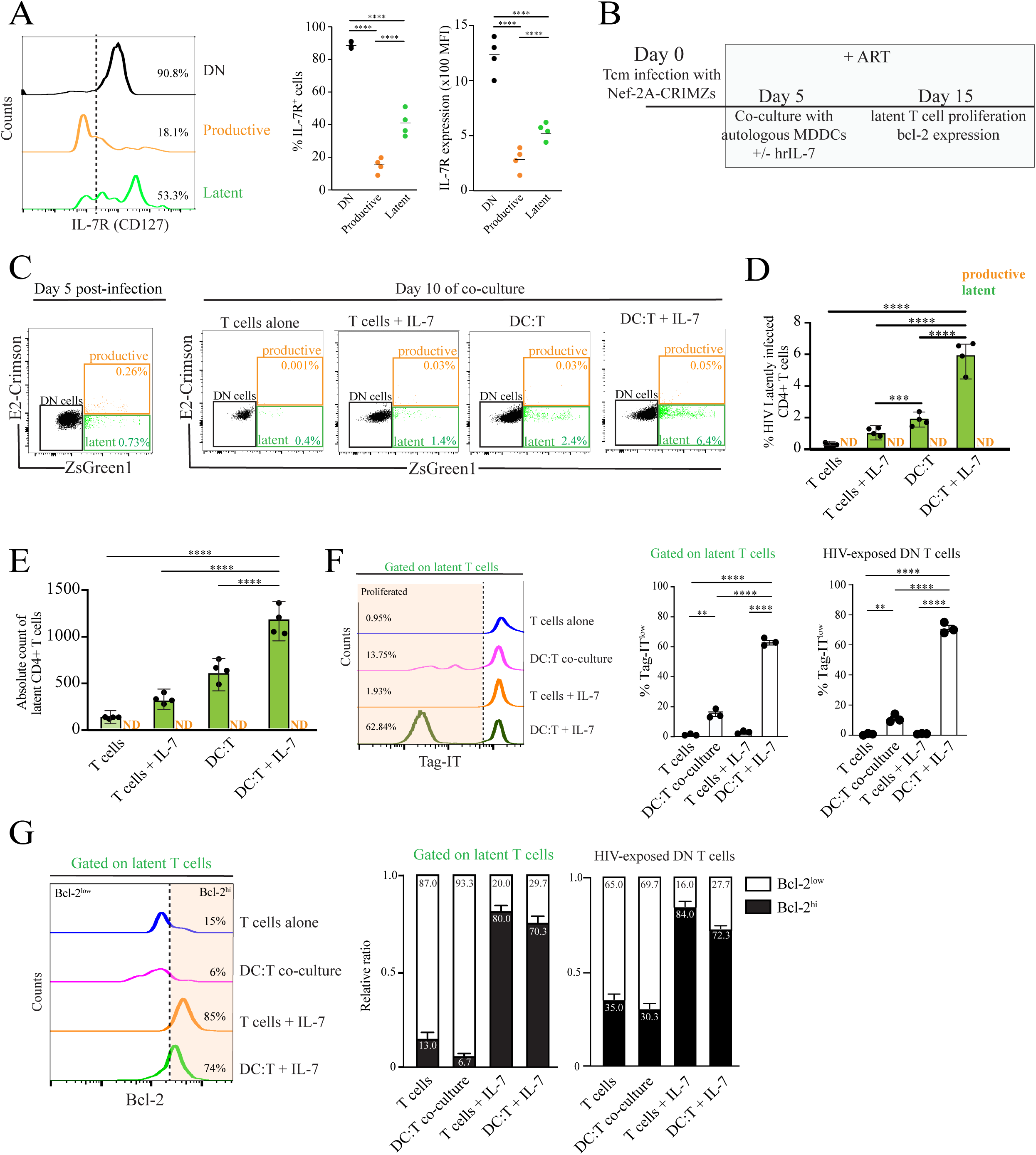
Presence of autologous dendritic cells and IL-7 promote a proliferative, pro-survival state in latent T cells. (**A**) IL-7 receptor expression was assessed in HIV-exposed, double negative (DN), productive and latent T cell populations at day 5 post-infection with HIV Nef-2A-CRIMZs. MFI = mean fluorescence intensity. **** p<0.0001, unpaired t-test. Representative data from 4 donors shown. **(B)** Experimental setup of autologous DC:T cell co-cultures in the presence of ART. Central memory-like T cells were stained with the Tag-IT proliferation dye and placed with autologous immature MDDCs at a ratio of 1:4 (T cell:DC) with raltegravir, emtricinabine and tenofovir. **(C)** Distribution of productive (orange box), latent (green box) and double-negative DN (black box) T cell populations by flow cytometry before and after 10 days in the indicated culture condition. **(D)** Flow cytometry analysis of latent and productive T cell population percentages after 10-day co-culture. ND = not detected. ****p<0.0001, Tukey’s multiple comparisons test. **(E)** Absolute cell counts of latent and productive T cell populations after 10-day co-culture. ND = not detected. ****p<0.0001, Tukey’s multiple comparisons test. **(F)** After gating on latent T cells, their proliferation based on Tag-IT dilution across different culture conditions are shown. Percent values indicate proportion of T cells that have undergone cell division relative to T cells alone (dotted line). Graphical representation of a representative donor out of 5 independent experiments is shown to the right, including HIV-exposed, DN T cells. **p<0.002, ****p<0.0001, Tukey’s multiple comparisons test. **(G)** Gating on latent T cells, Bcl-2 expression was assessed by intracellular flow cytometry. Percent values indicate proportion of T cells expressing Bcl-2 relative to T cells alone (dotted line; Bcl-2^hi^). Graphical representation of a representative donor out of 3 independent experiments is shown on the right.

### Cognate DC:T cell interactions drive proliferation and differentiation of virus-specific latent primary CD4^+^ T cells

Most of the long-lived latent HIV reservoir is contained within clonally expanded memory T cell populations in people living with HIV (PLWH) under ART, some of which are specific for viral antigens. We tested the hypothesis that cognate DC:T cell contacts are required for antigen-specific latent T cell proliferation in the absence of virus reactivation. We noticed that the addition of fetal calf serum (FCS), routinely used to supplement cell culture media, induced robust proliferation of latent and DN T cell subsets only in the presence of autologous DCs (Suppl. Fig. 3A-B). We interpreted these data as responses against bovine-derived antigens induced through DC:T cell contacts, rather than growth factors found in serum directly acting on T cell proliferation. These studies prompted us to perform all DC:T cell co-culture studies in defined media lacking FCS (Immunocult-XF) which supported high viability of both primary cell types, to reduce background T cell responses. Next, co-culture studies were performed in the presence of antigen pools containing viral lysates from Cytomegalovirus, Parainfluenza and Influenza virus (termed CPI lys) for up to 10 days. After infection of central-memory-like CD4^+^ T from healthy donors with HIV Nef-2A-CRIMZs, we show that the addition of DC+CPI lys helped promote robust expansion in the proportion and absolute cell counts of latent T cell populations (Fig. 3A,B). Similar responses were observed in HIV-exposed DN T cells in the same culture (Fig 3C), but no expansion of productively infected T cells was observed in our DC:T cell co-culture studies. To examine the role of dynamic DC and T cell interactions, we embedded DC:T cell co-cultures in either bovine or rat-tail 3D collagen matrices (both at 1.7mg/mL) in the presence or absence of CPI lys to evaluate latent T cell proliferative responses by flow cytometry (Fig. 3D). We and others showed that DC and T cell migration dynamics are drastically influenced by the two different 3D collagen matrix formulations: T cells in bovine collagen migrated at speeds comparable to those in the lymph node *in vivo*, whereas T cells in rat-tail collagen displayed very low motility due to smaller pore sizes that restrain cell movement (10, 46, 50, 51). No differences in T cell viability were observed up to day 5 in either matrix (Suppl. Fig. 4) but significantly higher levels of latent T cell proliferation was observed in bovine collagen compared to rat-tail collagen across 5 donors (Fig. 3D). These data suggest that constraints on T cell migration behaviors may prevent the formation of DC:T cell conjugates that are a requisite for antigen-driven proliferation. Notably, interactions with DCs led to T cell differentiation from a central memory to an effector memory-like phenotype with the latter population demonstrating higher proliferation levels compared to their T_CM_ counterparts (Fig 3E, Suppl. Fig. 5). Collectively, our data provide evidence that cognate DC:T cell interactions drive latent T cell proliferation in the absence of measurable HIV transcription.

**Figure 3:**
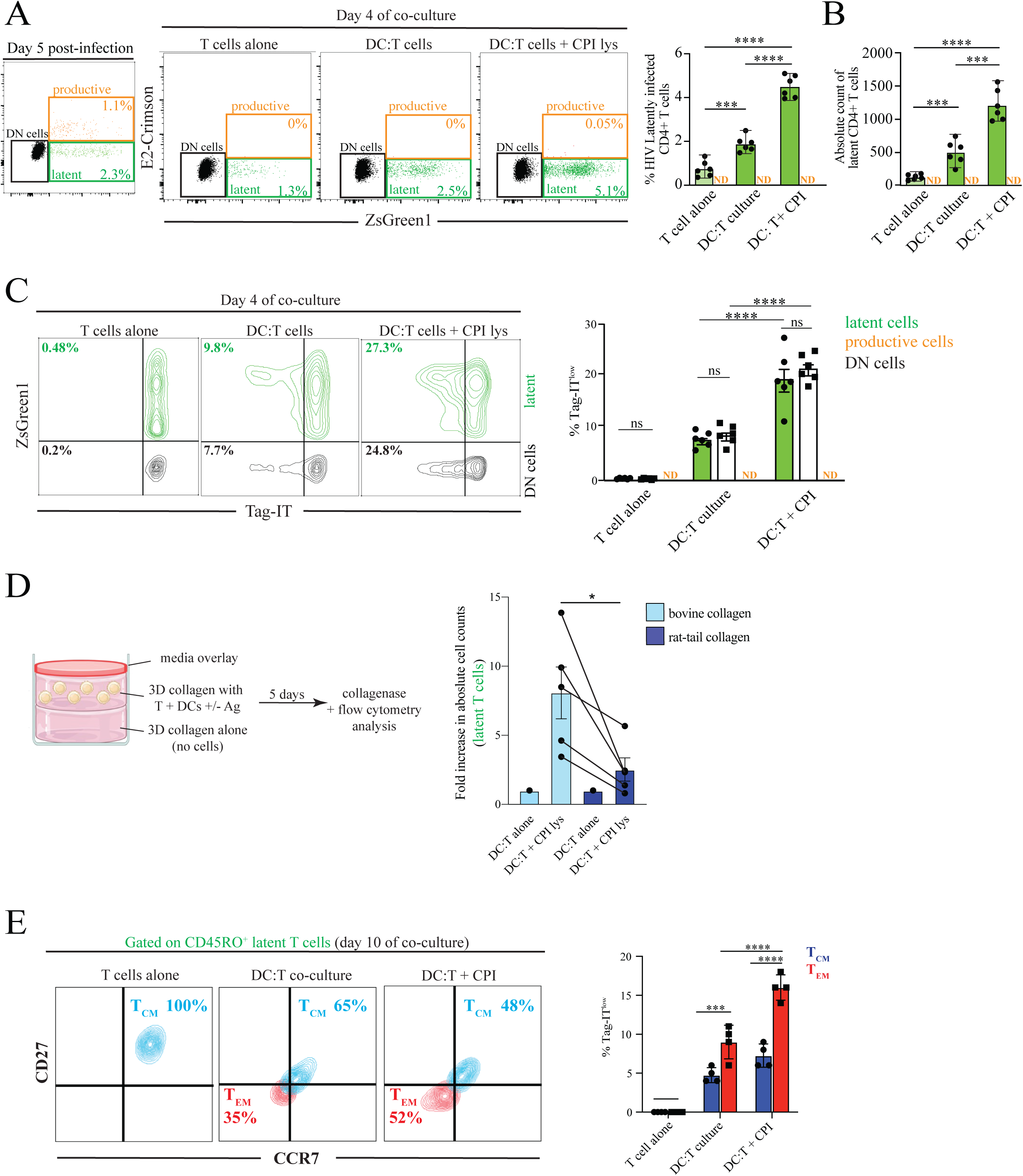
Cognate DC:T cell interactions drive virus-specific expansion of latent T cells. (**A**) T cells infected with HIV Nef-2A-CRIMZs were co-cultured with autologous DCs at a ratio of 4:1 in the presence or absence of pooled protein lysates from Human cytomegalovirus, Parainfluenza virus and Influenza virus (CPI lys). Distribution of productive (orange box) vs latent (green box) T cell populations by flow cytometry before and after 4 days of co-culture is shown. Graphical representation of a representative donor out of 6 independent donor experiments is shown on the right. ND = not detected. ***p<0.001, ****p<0.0001, Tukey’s multiple comparisons test. **(B)** Absolute cell counts of latent and productive T cell populations after 4-day co-culture. ND = not detected. ***p<0.001, ****p<0.0001, Tukey’s multiple comparisons test. Data from 6 independent donor experiments are shown. **(C)** Proliferative responses by Tag-IT dilution (4 days) across different culture conditions are shown. Both latent (green) and HIV-exposed double negative (DN; black) T cells from the same co-culture is shown. Percentages indicated cells that have divided based on Tag-IT dye dilution. Data from 5 independent donor experiments are shown to the right. ns = not significant, ****p<0.0001, Tukey’s multiple comparisons test. **(D)** DC:T cell co-culture in bovine or rat-tail collagen matrices in the presence or absence of CPI lys. Experimental design is depicted on the left. Right panel: fold increase in latent T cell numbers in co-cultures with CPI lys is shown for 5 donor cells. *p<0.05, Mann Whitney U test. **(E)** Phenotypic analysis of infected T cells in the presence or absence of autologous DCs (day 10) is shown. CD45RO^+^ memory T cells are gated and further stained for CCR7 and CD27. Percentages indicated for each compartment. Relative ratio of central and effector memory T cells is shown on the right. Representative data from 4 independent experiments is shown.

### Antigen presentation and co-stimulatory signaling drive clonal expansion of latent T cell pools

Initiation of antigen-specific T cell responses involve engagement with MHC-II and CD80/86 on DCs (16, 52–54). We next addressed whether these signals were also required for the expansion of recall T cell responses in HIV-infected cells. DC:T cell + CPI lys co-culture for 10 days led to proliferation (% Tag-IT^low^) and higher latent T cell numbers compared to T cell alone cultures, indicating active division over time across donors (Fig. 4A). We next show that MHC-II antibody blockade significantly reduced absolute cell counts and proportion of expanded latent T cell in all donors tested. Interestingly, CD80/CD86 blockade abrogated all T cell proliferative responses, including HIV-exposed, double negative T cell populations (Fig 4B,C). We confirmed receptor neutralization activity of all antibodies in DCs for up to 8 days by flow cytometry (Suppl. Fig 6). Additional controls using MHC-I antibody blockade showed no impact on latent T cell proliferation, confirming that the lack of responses was not due to non-specific disruption of DC:T cell contacts (Fig. 4D). DCs also upregulated MHC-II and CD80/86 expression during cognate DC:T cell interactions, likely through known positive feedback mechanisms including CD40/CD40L contacts (Suppl. Fig 7). We also stained for the anti-apoptotic molecule Birc-5 (Survivin) that is often upregulated in malignant cells (55, 56) and shown to prevent apoptosis of latently-infected CD4^+^ T cells (57). Birc-5 expression after 48 hours of DC:T cell co-culture was upregulated in the presence of CPI lys but abolished with CD80/CD86 blockade. Notably, Birc-5 expression was upregulated exclusively within dividing cells in both latent and HIV-exposed DN T cells (Fig 4E,F). Finally, we assessed the clonality of proliferating and non-proliferating latent CD4^+^ T cells by TCR/J sequencing of sorted T cell subsets. In contrast to non-proliferating T cells, high TCR clonality was observed within the proliferating T cell population (Fig 4G). Proliferating (Tag-IT^low^) and non-proliferating (Tag-IT^hi^) ZsGreen^+^CD4^hi^ T cells were sorted after 7 days of DC:T cell co-culture and processed for near full-length sequencing analysis of viral DNA. From the 20 amplicons we obtained sequences (8 non-proliferating, 12 proliferating ZsGreen^+^CD4^hi^ T cells), 1 intact provirus and 15 distinct viral DNA sequences were identified from our limited sampling, with some shared clones between the two populations and across cells from different donors (Fig 4H, Suppl. Fig 8). Together, cognate DC:T cell interactions drive antigen-specific expansion and pro-survival responses in infected T cells that is dependent on both TCR and co-stimulatory signals. Suppressing viral replication to restore normal T cell function may allow HIV-infected T cells to access physiological survival and proliferative cues in lymph nodes, ensuring long-term longevity of the viral reservoir in an antigen-driven manner.

**Figure 4:**
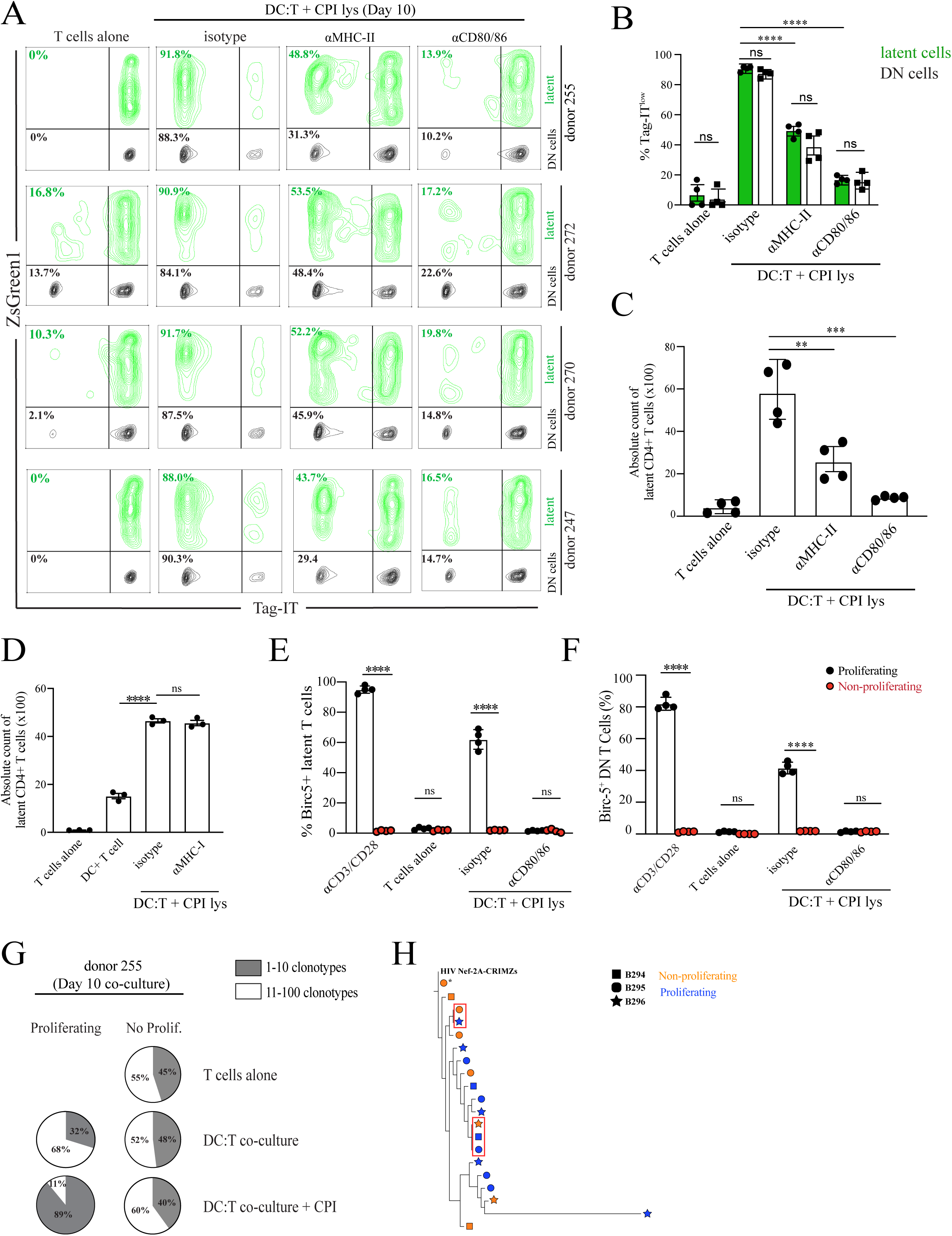
Antigen presentation and co-stimulatory signals drive clonal expansion of latent T cell pools. (A) Distribution of latent (green) and DN T cells (black) across 4 donors in the presence of either isotype or the indicated neutralizing antibodies. DCs were pretreated with blocking antibodies and were not removed throughout the duration of the DC:T cell culture. (B) % Tag-IT^low^ values across subsets and conditions indicated. Responses from 4 donors is shown. ns=not significant. **** p<0.001, Tukey’s multiple comparisons test. (C) Absolute cell counts of latent T cells after 10 days of co-culture with autologous DCs +/- CPI in the presence of either isotype or the indicated neutralizing antibodies. Responses from 4 donors is shown. (D) Cell counts of latent T cells in the presence or absence of MHC-I blocking antibody. Data is representative of 3 independent experiments. ns = not significant. **** p<0.0001, Tukey’s multiple comparisons test. (E) Birc-5 (survivin) expression was assessed in proliferating vs non-proliferating latent T cells in DC:T cell co-cultures. Representative data from 2 independent experiments is shown. ns = not significant. **** p<0.01, Sidak’s multiple comparisons test. (F) Birc-5 (survivin) expression was assessed in proliferating vs non-proliferating DN T cells in DC:T cell co-cultures. Representative data from 2 independent experiments is shown. ns = not significant. **** p<0.01, Sidak’s multiple comparisons test. (G) TCRb sequencing analysis of sorted T cells after DC:T cell co-culture from one representative donor. The proportion and frequency of clonal expansions in proliferating T cells is depicted. (H) Proliferating and non-proliferating CD4^+^ T cells were individually sorted for nested PCR amplification and near full-length sequencing. Phylogenetic trees for 3 donors (B294, B295, B296) built from the entire amplified area sequenced based on maximum likelihood. Sequences with 100% identify are boxed in red. * indicates intact provirus.

### Cognate DC:T cell derived cytokines have minimal effects on infected T cell proliferations

To further probe TCR-independent factors that can induce high proliferative responses in latent T cells, we performed multiplex cytokine arrays of clarified culture supernatants to quantitatively assess cytokine production. Using T cell cultures alone as baseline measurements, the presence of DCs on their own increased expression of several cytokines including TNF, IL-9, IL-15, IL-6 and IL-12p40, but expression was significantly enhanced in the presence of CPI antigen pools (Fig 5A). Reduced secretion of cytokines after MHC-II and CD80/86 blockade was observed, indicating that direct DC:T cell contacts were a requirement for cytokine release. DC:T cell culture in the presence of the superantigen staphylococcus enterotoxin B (SEB) was used as a positive control. Of note, IL-7 was not detected in the culture media in all conditions. Next, culture media from various conditions in Fig. 5A were applied to latent T cell populations generated from the same donor for 10 days. When compared to the robust proliferative responses observed in the presence of DCs+CPI lys, culture media alone had minimal effects on T cell proliferation (Fig 5B-C). We conclude that while antigen-driven DC:T cell contacts induce production of cytokines that may have the capacity to stimulate T cell proliferation, they had little contribution on HIV-infected T cells expansion on their own.

**Figure 5:**
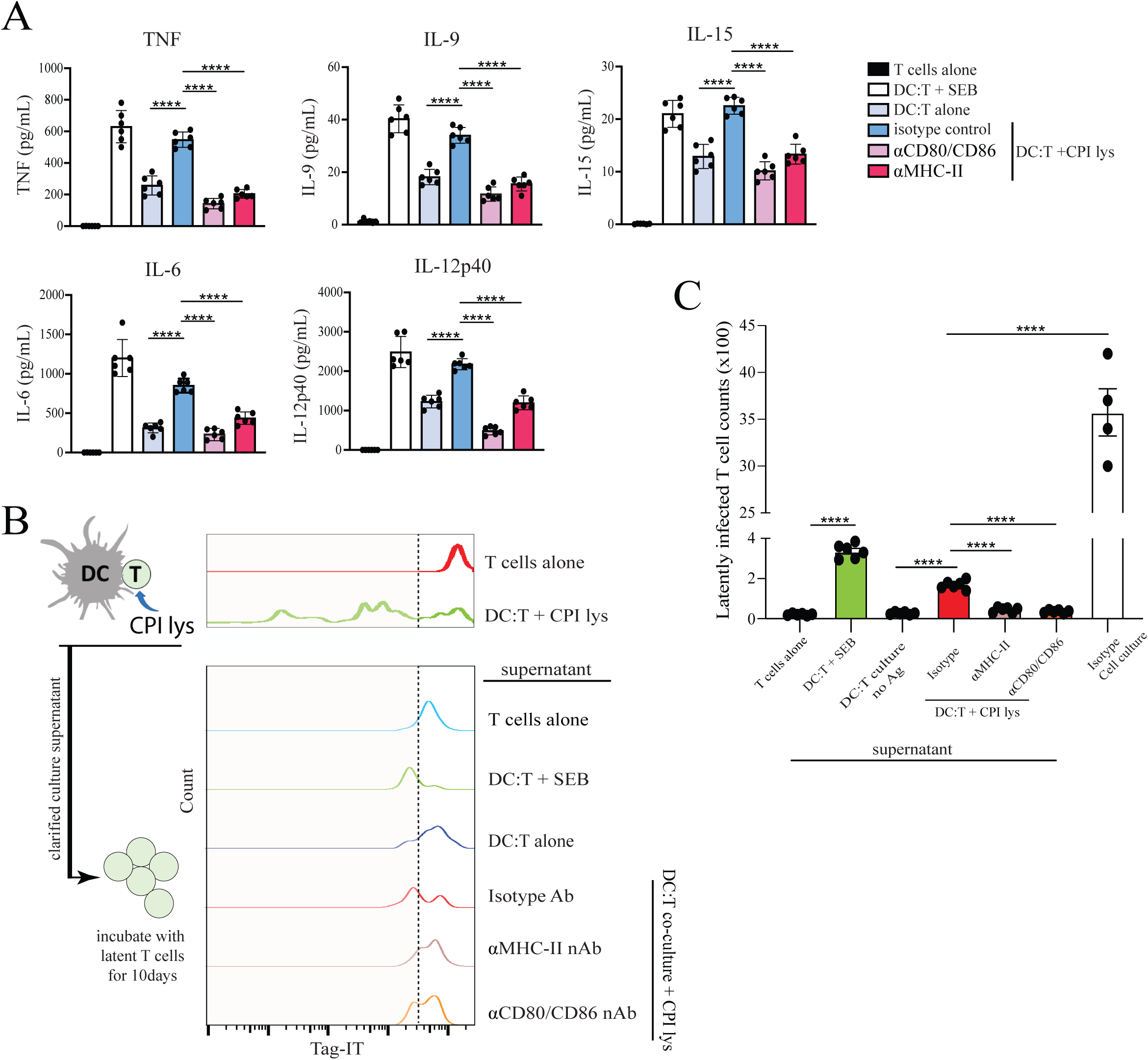
Cognate DC:T cell derived cytokines have minimal effects on latent T cell proliferation. (**A**) Culture supernatant from the indicated conditions were harvested at day 10 and assessed by multiplex ELISA for TNF, IL-9, IL-15, IL-6 and IL-12 expression. Representative data from 6 independent experiments is shown. **** p<0.0001, Tukey’s multiple comparisons test. SEB = staphylococcus enterotoxin B. **(B)** Latently infected T cells were cultured with media supernatant collected from the same donor in (A) and their proliferative responses assessed after 10 days. Proliferative responses by T cells in DC:T cell co-cultures are included at the top of the figure panel for response comparison. Representative data from 6 independent experiments is shown. **(C)** Absolute cell counts after incubation with culture supernatants. While some level of proliferation was observed with culture media, proliferative responses were low compared to DC:T cell co-cultures + antigen. Representative data from 6 independent experiments is shown. ****p<0.0001, Tukey’s multiple comparisons test.

### Modulating TCR signaling strength uncouples HIV reactivation from proliferative responses in infected T cells

T cell activation and HIV transcription are often viewed as an interconnected process downstream of the TCR, but recent studies show that these responses can occur independently of one another (4, 30, 58, 59). We postulated that graded TCR signals were a key regulatory factor that controlled a range of downstream responses in HIV-infected T cells, with the induction of proliferative responses in the absence of overt proviral reactivation favoring long-term HIV persistence. When latent T cells were stimulated with increasing amounts of anti-CD3/CD28 conjugated beads, we observed that a minimum threshold stimulus was required to induce cell proliferation with little to no Gag p24 expression (T cell:bead ratio of 1:0.5; Suppl. Fig. 9A). CD69 expression predictably increased with higher amounts of stimulating beads in this assay (Suppl. Fig. 9B). Similar observations were made when DC:T cell ratios were varied with a fixed amount of CPI lysis antigen (Suppl. Fig.9C-D) consistent with a previous study (60). In both instances, low levels of TCR stimulation were sufficient to drive T cell proliferation with minimal Gag p24 re-expression, whereas incremental increase in TCR signaling (i.e., high bead:T cell or DC:T cell ratios) enhanced Gag p24 re-expression potential (Suppl. Fig. 9C) with a corresponding reduction in proliferative responses possibly due to viral cytopathic effects (Suppl. Fig 9E).

The limitation of these experimental approaches is that individual TCR signaling events cannot be directly attributed to a specific T cell response within a polyclonal population. To address this, we utilized two primary CD4^+^ T cell clones derived from SARS-CoV-2 infected individuals (clone P155/P156) that expressed high GrzB and IFNγ levels upon cognate peptide stimulation presented in the context of autologous B cells (61). When P155 and P156 clones were infected with HIV Nef-2A-CRIMZs, similar infection rates and latent T cell phenotypes to polyclonal T cells were observed at day 5. Infected T cell clones were co-cultured with autologous B cells with increasing cognate peptide concentrations and phenotypic analysis were determined by flow cytometry (Fig. 6A). Baseline expression of HLA-DR and PD-1 were low but increased with peptide stimulation in a dose dependent manner, as expected (Fig. 6B,C). At 8 days of T:B co-culture, up to >60% of ZsGreen^+^CD4^hi^ T cells underwent cell division at the 50-100ng peptide dose (Fig 6D), and this was confirmed by increase in absolute cell counts (Fig 6E). Increasing peptide dose in this system led to a graded increase in Gag p24 expression that corresponded with a significant reduction in latent T cell numbers at the 400ng and 1μg peptide concentrations (Fig 6D). Notably, Gag p24^+^ cells expressed high levels of HLA-DR compared to their p24^-^ counterparts (Fig 6D). These data support the notion that weak TCR stimuli (which is sufficient to induce modest levels of PD-1 expression) can induce proliferation of infected T cells with minimal proviral reactivation. We also observed that latent T cells expressed higher PD-1 levels (Fig 6D), and that PD-1 blockade in T:B co-cultures at the 100ng peptide dose resulted in higher T cell activation (HLA-DR^+^), increased Gag p24 re-expression and lower numbers of expanded latent T cells (black bars) (Fig 6F-H). These studies suggest that the removal of negative regulatory signals increases TCR signaling magnitude that increases latency reversal resulting in reduced proliferative responses (Fig 6E-F). These observations argue that the magnitude of antigenic TCR stimulation is likely a key determinant that drive clonal expansion of select HIV-infected but transcriptionally silenced T cell subsets, some of which dominate the persistent HIV reservoir over years on suppressive ART.

**Figure 6:**
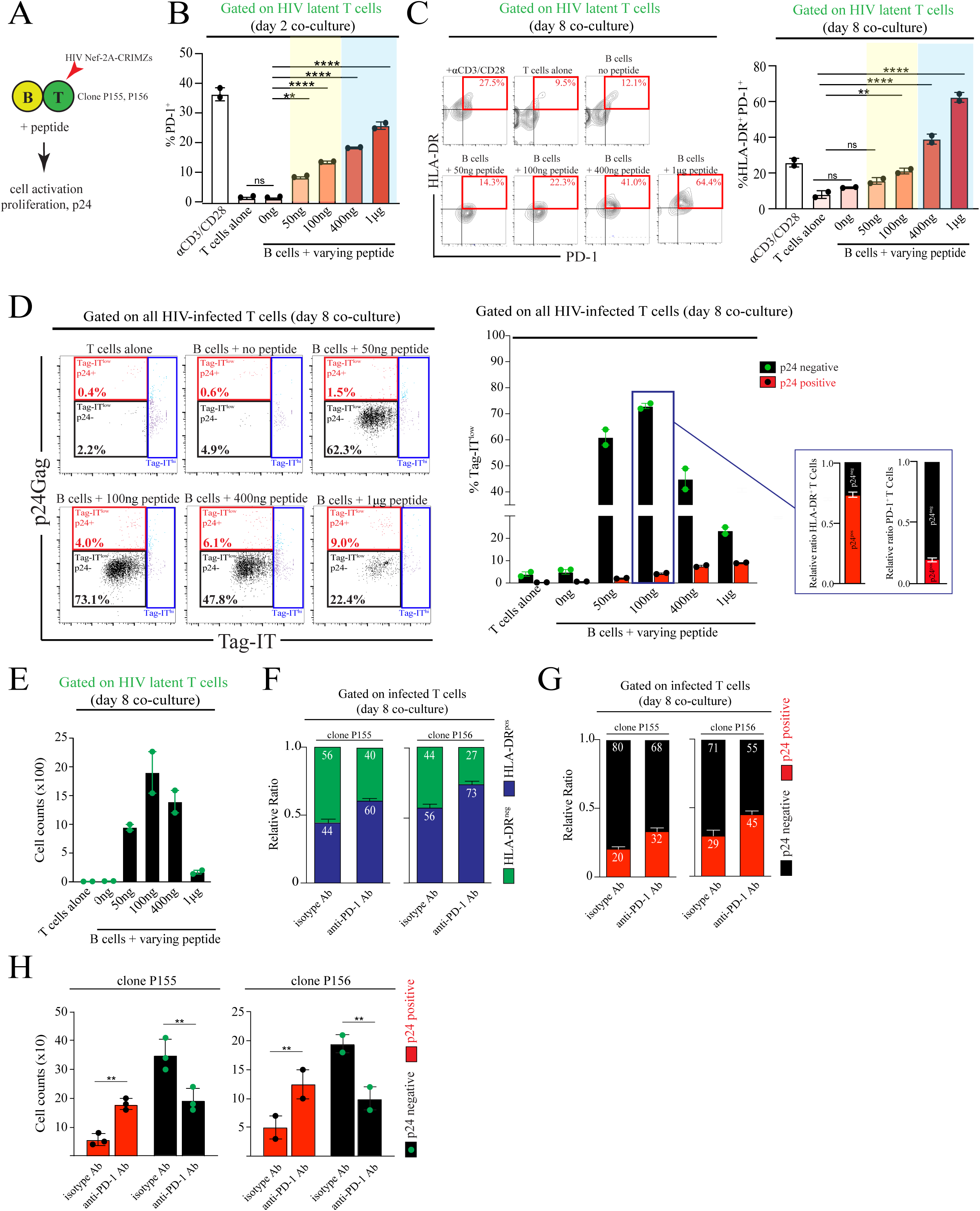
Graded TCR signaling uncouples T cell activation, cell proliferation and HIV reactivation potential in clonal T cells. (A) SARS-CoV-2 specific CD4^+^ T cell clones P155 and P156 were infected with HIV Nef-2A-CRIMZs and co-cultured with autologous B cells in the presence or absence of cognate peptide. T cell activation, proliferation and Gag p24 expression was evaluated by flow cytometry. (B) Gated on latent T cells, the proportion of T cells expressing PD-1 and HLA-DR in the presence of increasing peptide concentrations (ng/mL) was evaluated at 2 and 8 days of T:B co-culture, respectively using flow cytometry. Representative data using clone P155 is shown. ns = not significant. **p<0.002, ****p<0.0001, Tukey’s multiple comparisons test. (C) Flow plot of HLA-DR and PD-1 expression across conditions using clone P155. %HLA-DR+PD-1+ T cells is shown to the right. ns=not significant. **p<0.002, ****p<0.0001, Tukey’s multiple comparisons test. (D) Proliferative response using Tag-IT dilution was measured in all infected T cell clone P155. Responses were further divided to T cells expressing or not expressing Gag p24. HLA-DR and PD-1 expression in p24^+^ and p24^neg^ cells are also shown. (E) Absolute cell counts of latent T cells across different T:B co-culture conditions. (F) Relative ratio T cells expressing HLA-DR in T:B co-cultures with or without PD-1 blockade. All HIV-infected T cells are gated in this analysis. Data using clones P155 and P156 are shown. (G) Relative ratio T cells re-expressing Gag p24 in T:B co-cultures with or without PD-1 blockade. All HIV-infected T cells are gated in this analysis. Data using clones P155 and P156 are shown. (H) Absolute cell counts in T:B co-cultures with or without PD-1 blockade. **p<0.01, Sidak’s multiple comparisons test.

## Discussion

HIV persistence despite years of ART suppression poses a major barrier to achieving HIV cure in PLWH (2, 4, 62, 63). An outstanding question is whether the cellular and molecular signals that support memory T cell homeostasis contribute to the maintenance of the T cell reservoir under ART. In this study, we used a fluorescent reporter-based approach to assess whether DC-derived cues directly modulate the activation potential and proliferative capacity of HIV-infected, transcriptionally silent (ZsGreen^+^CD4^hi^) CD4^+^ T cells in a series of *ex vivo* DC:T cell co-cultures studies. We show that cognate DC:T cell interactions drive clonal expansion of ZsGreen^+^CD4^hi^ T cell subsets, and that a pro-survival state within proliferating cells is reinforced through IL-7 signaling. Interestingly, we describe a dominant role for CD28 co-stimulation in regulating robust latent T cell proliferation which was associated with upregulation of anti-apoptotic Birc-5 expression. Using primary CD4^+^ T cell clones, we show that an incremental reduction in TCR stimulation initiated robust proliferative responses in the absence of measurable viral reactivation, whereas saturating TCR stimulation increased HIV transcription and lowered their expansion potential. PD-1 blockade during cognate interactions resulted in higher TCR signaling, increased HIV Gag expression and reduced proliferative capacity, leading to an overall reduction in the number of expanded latent T cells. Collectively, our studies argue that the magnitude of TCR/co-stimulatory signaling during cognate APC:T cell interactions are key regulators of the underlying proliferative and survival programs maintaining reservoir persistence. Inducing a latent state may allow infected T cells to resume physiological behaviors and functions *in vivo*, some of which contribute directly to the maintenance of the HIV reservoir.

Longitudinal, single-cell analysis of HIV-infected CD4^+^ T cell pools indicate that the latent reservoir *in vivo* is largely maintained through clonal proliferation and shaped by epigenetic regulation and immunological control that favor their long-term persistence (64, 65). Several mechanisms that help explain proliferative responses of infected cells *in vivo* have been described. Proviral integration into cancer related genes such as *STAT5B* can promote cell-autonomous proliferation, although this mechanism only partially describes the clonal nature of the latent reservoir (25, 35, 65, 66). Similarly, the importance of homeostatic cytokines such as IL-7 and IL-15 on reservoir maintenance is well documented (15, 45, 67) but are unlikely to exclusively drive expansion of specific T cell clones over time. Several groups have identified clonally-expanded T cells harboring replication competent provirus with antigen specificities to viral (Epstein-Barr virus, CMV, HIV Gag, and Influenza virus), bacterial (*S. aureus*) and fungal-derived (*C. albicans*) pathogens that were not explained by homeostatic proliferation or specific integration site effects (24, 25, 28, 29). These studies indicate that continual exposure to microbial antigens may drive reservoir persistence by stimulating proliferative and pro-survival programs in infected T cells, thereby offsetting cell loss through viral cytopathic effects or ongoing immune regulation. A recent study also showed that presence of self-derived antigens (from non-B and T cell lysates) can stimulate viral production in individuals experiencing nonsuppressible viremia in an MHC-II dependent manner (26). Observations that viral blips are more common during winter seasons (68) and after flu immunization (69) reiterates the possibility that T cell responses are continually regulated by microbial and self-derived stimuli in PLWH, which in some instances may favor the expansion of specific T cell clones harboring provirus. In this study, we sought to better understand the molecular cues that favored HIV persistence through antigen-driven proliferation.

Proviral sequencing analysis of tissue samples obtained from PLWH donors on suppressive ART post-mortem reveals high clonality of the HIV reservoir across a wide range of organ systems (highest in secondary lymphoid organs), suggestive of extensive recirculation of HIV-infected cells and their local expansion within tissues (22, 23). While these studies did not distinguish the cellular reservoirs at these anatomical sites, we showed previously that HIV-infected CD4^+^ T cells retain robust motility within lymph nodes and facilitate viral dissemination through active migration into distant lymphoid organs (5–7). Migrating T cells in SLOs engage with a stable network of DCs along fibroblastic reticular cells (FRCs) that, upon cognate antigen recognition, results in T cell activation, proliferation and differentiation over the next few days (70, 71). In the absence of antigen, T cells actively leave the lymph node via cortical sinuses in a S_1_PR1-mediated, multistep process and enter other lymph nodes (72). Our studies show that while tonic DC:T cell interactions can support low level proliferation, the presence of cognate antigen greatly enhances clonal proliferation and induces a pro-survival program within responding cells. Importantly, we confirm that absolute infected T cell counts significantly increase in the presence of DC+antigen which indicates active cell division over time, both in suspension and in a 3D collagen matrix environment. More recent studies show that CD4^+^ T cells harboring HIV provirus in ART suppressed individuals display transcriptional and epigenetic signatures associated with a strong proliferative program (*IKZF3*, *BIRC5, BACH2* among others) and inhibition of death receptor/necroptosis pathways (36, 37, 39, 73). This is consistent with studies by Kuo *et al* showing that Birc-5 upregulation in latent T cells enhances their survival (57), and that bcl-2 upregulation in HIV-infected T cell pools increased resistance to CTL killing (74). We extended these findings by showing that Birc-5 expression was upregulated exclusively within proliferating T cells upon cognate DC:T cell interactions. While IL-7 has been shown to induce low proliferative responses without viral reactivation (45, 58), this was not observed in our assays but IL-7 did upregulate bcl-2 expression, a known downstream effector molecule of the IL-7 signaling pathway (75, 76). Combined with findings that lymph node homing chemokines CCL19/21 enhance HIV latency in proliferating and non-proliferating T cells (77) and observations that APCs can directly enhance latent infections in a cell-cell contact dependent manner (78–80), our studies highlight the role of lymphoid tissues as central hubs that support the generation and maintenance of the latent reservoir, a model that is supported by *in vivo* data (23, 31, 81–83). We propose that the secondary lymphoid tissue architecture help promote antigen-specific expansion of infected T cell clones under ART by facilitating rare cognate DC:T cell interactions to occur in the presence of homeostatic cytokines and chemokines (84).

TCR engagement with the peptide-MHC complex was required for clonal proliferation of virus-specific, HIV-infected T cell subsets in our studies, with minimal contributions by secreted cytokines alone. We observed a strong requirement for CD28 co-stimulation in this process that was consistent across all donors tested. It is well established that induction of primary T cell responses requires both cognate TCR:pMHC interactions and CD28 co-stimulation (85–88) and a long-standing notion that optimal secondary T cell responses do not require CD28 inputs. However, accumulating studies have challenged this concept in both mouse (89, 90) and humans (91, 92) and collectively show that CD4^+^ and CD8^+^ memory T cells require CD28 signaling for maximal expansion and pathogen clearance (93). Our observations are in agreement with these studies and show that CD28 signaling play crucial roles in the expansion of memory T cells by linking to proliferative, survival and metabolic programs regulated by the downstream PI3K/AKT pathway. Cognate DC:T cell contacts induced higher proliferation in effector memory T cells which aligns with studies reporting a higher proportion of inducible, replication-competent provirus in effector memory T cell subsets (83, 94). However, other studies reported similar distribution of intact provirus across memory subsets, and repeated stimulation led to differentiation towards an effector phenotype which could also explain our observations (95). Recent studies demonstrated that T cells with intact provirus displayed reduced cell proliferation compared to cells harboring defective provirus, which intriguingly was not associated with higher virus production after stimulation with anti-CD3/CD28 conjugated beads (59). In our limited sampling of sorted proliferating and non-proliferating infected T cells, 1 cell with intact provirus was detected in non-proliferating, but not in expanded populations, whereas small clones harboring defective provirus were observed after DC:T cell co-culture. While our observations seem to align with these studies, our sample size was too low to make broad conclusions regarding the proliferative capacity of T cells containing intact versus defective provirus. Contribution of integrins (eg LFA-1) and other costimulatory molecules (eg OX40, 4-1BB) in modulating cell-cell interaction dynamics and fine-tuning proviral reactivation versus proliferative signals in latent T cells harboring intact vs defective provirus is an important question and subject of future investigations.

Studies with human CD4^+^ T cell clones allowed us to dissociate T cell-intrinsic from extrinsic factors that regulate cellular activation, HIV transcription and proliferative responses. The overall strength of the TCR signal is a sum of TCR affinity, pMHC-II densities and costimulatory signals, each influenced by the dynamic evolution of chronic infections. Stimulation with decreasing peptide concentrations, while keeping the number of APCs constant, induced modest HLA-DR and PD-1 expression but was associated with robust proliferative responses in the absence of measurable Gag p24 expression. Increasing peptide dose resulted in higher activation, notable increases in HIV transcription and significant reductions in absolute latent T cell numbers after 8-10 days of co-culture. These data argue that the minimum threshold signal required to induce proliferative responses is lower than those required for proviral reactivation, and that the uncoupling of these two responses likely favors HIV persistence. Findings that a large fraction of intact provirus was not induced even after four rounds of T cell stimulation supports this hypothesis and indicate that proviral inducibility is restrained by epigenetic regulation, integrational constraints, and availability of key transcription factors (65, 95). Thus, proviral reactivation on a per cell basis is modulated by the strength and nature of the T cell stimulus and specific proviral silencing mechanisms in place. Studies using chimeric antigen receptors (CARs) of varying ligand binding affinities showed that receptors with low and intermediate binding affinities failed to support viral transcription, whereas high affinity CARs did (96). We further probed this concept in our model system by disrupting PD-1 signaling, which primarily inhibits TCR-linked signals (97). PD-1 blockade during cognate T:B cell interactions resulted in higher TCR signaling potential, increased Gag p24 re-expression and a corresponding reduction in proliferative capacity, possibly due to viral cytopathic effects. Interestingly, we observe that some Gag p24^+^ cells maintain cell division, which is consistent with clonal expansion of T cells displaying strong proviral transcriptional activity in ART-suppressed individuals and contribute to the latent reservoir (65). Immune checkpoint (IC) molecules LAG-3, PD-1 and TIGIT are preferentially expressed by persistently-infected CD4^+^ T cells and PD-1 signaling can reduce T cell activation and directly silence HIV transcription (98–101). Single-cell phenotypic analysis confirmed PD-1 and TIGIT expression in latent T cells which play a role in enhanced survival (64), resistance to cell-mediated killing (101, 102) and PD-1 blockade has shown clinical promise as a latency reversing agent (103). Recent studies show that ongoing HIV-specific T cell responses are maintained by spontaneous proviral transcription, and that cells harboring transcriptionally active provirus are progressively eliminated by host immunity (65, 104). In our co-culture systems, productively infected T cells were rapidly lost over time, likely due to cell death and only those that remained transcriptionally silent continued to survive *in vitro*. We speculate that high affinity clones are initial sources of stochastic viral transcription in response to infection (eg influenza A virus) that are progressively purged by cell-mediated immune responses, whereas the remaining pool of lower affinity clones remain in a latent state and eventually dominate the latent reservoir through antigen-driven expansion. This concept is reminiscent of findings that lower affinity/avidity clones of responding T cells dominate during late stages of pathogen-specific responses *in vivo* after high affinity T cells experience Activation-Induced Cell Death (AICD) or immune exhaustion (105–107). We acknowledge that our approach does not assess the proportion of latent T cells that reactivate virus production and die upon cognate DC:T cell interactions and is subject of our ongoing investigations. Nevertheless, our studies suggest that the overall strength of the TCR signal accumulated during blood and tissue recirculation is another important factor that regulate which HIV-infected T cell clones ‘wax and wane’ during long-term ART. HIV-infected T cells could be stimulated to proliferate, and in the absence of proviral reactivation, the cells could avoid viral cytopathic effects and immune clearance.

This study has several limitations. We note that any model system used to generate an HIV latent state in culture is not subject to epigenetic and metabolic changes imposed by host immunity observed under long-term ART. However, our validated latency reporter construct allows for confirmational, mechanistic studies into the molecular regulation of HIV latency, and the behavior and cellular responses to stimulus which are studies that are difficult to perform using patient-derived cells. We argue that studies isolating the contribution of TCR-derived signaling (using CD4^+^ TCR clones for example) is also not possible without a reporter-based HIV infection construct. We observe robust antigen-driven proliferation of latent T cell subsets that is dependent on cognate APC:T cell interactions, but clonality cannot be definitively established without additional integration site analysis. However, we noted minimal to no latent T cell proliferation in the absence of autologous DCs and CPI antigen pools, suggesting that the site of proviral integration has little effects on proliferative responses observed in our study. HIV infection studies using a similar primary central memory CD4^+^ T cell model reported that ∼50% of infected T cells contained intact provirus, with the majority remaining in a non-inducible state after TCR stimulation (108). In our limited sample size, we observed that most proliferating T cells contained defective provirus in *Env*, *Nef* and *Crimson* genes: Thus, sequence analysis of a larger number of proviruses is needed for more broad conclusions regarding the proliferative capacity of T cells containing intact versus defective provirus in our DC:T cell co-cultures. Finally, PBMCs were isolated from anonymized blood donors with no information on sex/gender, age or health status at the time of this study and thus the contribution of sex on T cell responses was not addressed. Segregated analysis based on sex, in conjunction with information on recent infections, are important considerations and will be addressed in future studies to define factors that impacts the breadth and magnitude of latent T cell responses.

In conclusion, we developed a fluorescent HIV latency reporter system to define the molecular underpinnings that drive antigen-specific expansion of HIV infected T cell pools through cognate cell-cell contacts. We show that HIV-infected but transcriptionally silent T cells retain their ability to engage APCs in an antigen specific manner, thereby driving a proliferative and pro- survival program that is similar to physiological T cell responses. Our model system enables defining how the peptide affinity range impacts the extent of T cell latency expansion and provides a rationale for an immunological-based therapeutic approach to reducing reservoir size in PLWH.

## Materials and Methods

### Cells

Human CD4^+^ T cells and MDDCs were isolated from PBMCs of donors (NetCAD, Canadian Blood Services). All cells derived from donors are anonymized and contain no information regarding the sex/gender, race, age or health status. All studies with human blood products were approved by the University of Manitoba Biomedical Research Ethics Board (B2015:030) and NetCAD (CBSREB 2020-018).

MAGI.CCR5 cells were obtained from the NIH HIV Reagents Program (Cat #3522) and grown in DMEM supplemented with 10% fetal calf serum (VWR Seradigm), 2 mM GlutaMAX (Gibco), 1mM sodium pyruvate (Corning) and 10mM HEPES (Sigma-Aldrich) under 37°C/5% CO_2_ conditions. The parental cell line of MAGI is a HeLa cell clone, which is a female cell line. This cell line has not been authenticated.

### Construction and validation of the dual-fluorescent HIV reporters

The E2-Crimson-EF1a-ZsGreen DNA insert was amplified from the Hi.Fate latency plasmid (109) using primers 5’-TGC ACG CGT GGA GGG GGC GGT ATG GAT AGC ACT GAG AAC G-3’ and 5’-GCT ACC CGG GTC AGG GCA AGG CGG AGC CGG AGG CG-3’ by PCR, and inserted into the R5-tropic ‘HIV-GFP’ proviral vector (5) using unique restriction enzyme sites XmaI (NEB # R0180S) and MluI (NEB # R0198S). The P2A sequence from Porcine Teschovirus-1 (110) was generated using custom oligonucleotides and inserted into the MluI site between the *Nef* and *E2-Crimson* gene locus. The resulting plasmid, termed ‘HIV Nef-2A-CRIMZs’, was sequenced on both strands before transfection into HEK293T cells. To construct the HIV Nef-ZTom reporter, the dTomato gene amplified using primers 5’-GCC ACC ATG GTG AGC AAG GGC GAG GAG GTC-3,’ and 5’-GCT ACC CGG GTT ACT TGT ACA GCT CGT CCA-3’ by PCR and inserted into the R5-tropic HIV Nef-2A-CRIMZs plasmid using XmaI and NcoI sites (this replaces the original Zsgreen1 with dTomato gene). The Nef-ZsGreen fusion gene was generated previously and amplified using primers 5’-TGC ACG CGT GGA GGG GGC GGT ATG GCC CAG TCC AAG CACG-3’ and 5’-AGA TCC GCG GTC AGG GCA AGG CGG AGC CGG-3’ by PCR. Viral supernatant was collected and concentrated, as described below.

### Viral Constructs and Preparation of Viral Stocks

All HIV stocks were produced by transient transfection of HEK293T cells (ATCC; CRL-3216) using JetOPTIMUS solution. In some cases, HIV stocks were VSVg pseudo-typed to achieve high infection rates. All viral supernatants were harvested at 48 hours post-transfection and centrifuged at 500 xg for 10 mins to remove cellular debris. Culture supernatant was mixed with Lenti-X concentrator (Takara Bio; cat # 631232) for 60 mins prior to centrifugation at 1500 xg for 45 mins at 0°C. Viral pellets were resuspended in 1/100 original volume of complete DMEM (100X concentration) and viral stock aliquots stored at -80°C. All viral stocks were titrated using MAGI.CCR5 cell lines and expressed as blue-forming units (bfu) per milliliter. In all experiments, CD4^+^ T cells were infected with HIV Nef-2A-CRIMZs at a multiplicity of infection (MOI) of 0.1.

### Isolation and preparation of primary cells

CD4^+^ T cells and CD14^+^ monocytes were isolated from PBMCs of anonymized human donors using the EasySep Human Naïve CD4+ T cell Isolation Kit II (Stemcell Technologies; cat #17555) following manufacturer protocols with cell purity routinely >95%. MAGI.CCR5 cells were obtained from the NIH AIDS Reagents Program (#3522) and cultured in RPMI supplemented with 10% fetal bovine serum (VWR Seradigm), 1mM sodium pyruvate (Corning; cat #25-000-CI), 2 mM GlutaMAX (Gibco; cat #35050061) and 10mM HEPES (Sigma-Aldrich; cat #1563008) under 37°C/5% CO_2_ conditions. The parental cell line of MAGI.CCR5 is a HeLa cell clone, which is a female cell line. This cell line has not been authenticated. HEK293T cells were obtained from ATCC (#3522) and cultured in DMEM supplemented with 10% fetal bovine serum, 1mM sodium pyruvate, 2mM GlutaMax and 10mM HEPES under 37°C/5% CO_2_ conditions.

### Cell culture

Monocyte-derived dendritic cells (MDDCs) were generated by isolating CD14^+^ monocytes and seeding into T25 NunclonTM Sphera flasks (Thermo Scientific; cat #12566439) in Immunocult-XF cell expansion media (Stemcell Technologies; cat #10981) supplemented with 1% Penicillin/Streptomycin and 50ng/mL each of human granulocyte macrophage colony-stimulating factor (hGM-CSF; Biolegend, cat #578606) and interleukin 4 (Biolegend; cat #574008) for five days. Flow cytometry and morphological analysis confirmed differentiation in MDDCs (10). To generate autologous central memory T cell populations, naïve CD4^+^ T cells from the same donor were activated with anti-human CD3e/CD28 antibody coated Dynabeads (1:1 bead:cell ratio, Life Technologies; cat #11131D) and expanded in the presence of rhTGFb (10ng/ml), anti-human IL-4 (1μg/ml), anti-human IL-12 antibody (2μg/ml) and anti-human IL-23 antibody (2μg/ml) in Immunocult-XF expansion media, as previously described (111). After 48 hours, beads were magnetically removed using an EasySep magnet (Stemcell Technologies; cat# 18000) and T cells were cultured for another 5 days in Immunocult-XF media supplemented with 50 IU/mL human recombinant IL-2 (R&D systems; cat #200-02), keeping cell densities at 2 x 10^5^ cells/mL. Flow cytometry analysis confirmed T_CM_ (CCR7^+^CD27^+^) and resting (HLA-DR^low^/CD69^low^) phenotypes. Day 7 expanded T cells and day 5 differentiated MDDCs were used for all co-culture studies.

### Flow cytometry and antibodies

CD4^+^ T cells and MDDCs were stained for their respective surface makers using conjugated antibodies (STAR Methods). Briefly, DCs were first pretreated with 1μL of Fc receptors (FcRs) blockade using human TruStain FcX solution (Biolegend; cat #422302), and incubated on ice for 5 mins. Afterwards, cells were stained with the indicated surface antibodies for 30 minutes. Cells were washed twice by adding 2 mL of FACS buffer (PBS + 2% FCS) and centrifuged at 400 xg for 5 mins. The supernatant was discarded, and cells were resuspended in 500 μL of 2% PFA. Cell acquisition was performed on the FACS Canto II (BD Bioscience, USA). Data was analyzed using FlowJo software (version 10.7.2). In some studies, infected T cells were stained with anti-CD3 and CD4 antibodies and live sorted for ZsGreen^+^CD3^+^CD4^high^ latent T cells using the BD FACS AriaFusion. To assess HIV reactivation potential, sorted cells were stimulated with anti-human CD3/CD28 activating beads (StemCell Technologies; cat #10971) for 24 hours, then assessed for p24Gag and HLA-DR/CD69 expression by flow cytometry.

### HIV infection of T cells in 3D collagen matrix

T cells were infected in 3D collagen matrix, as previously described (46). Briefly, 2 x10^7^ day 7 expanded CD4^+^ T cells were pelleted in a FACS tube by centrifugation at 400xg for 6 minutes, then infected with 0.1 MOI HIV Nef-2A-CRIMZs in bovine collagen solution at a final concentration of 1.6 mg/mL collagen in a 48-well plate for 3 hours. After the incubation period, 400μL of 1x collagenase D (Roche; cat #11088866001/ final concentration of 1mg/mL) was added for 15 minutes to remove T cells, washed and resuspended at 10^6^ cells/ml in Immunocult expansion media, supplemented with rhIL-2 (50IU/mL) +ART (more detail below) in a 24-well plate. Media with IL-2+ART was replaced every 2 days. Flow cytometry at day 5 post-infection confirmed infection for downstream co-culture studies.

For DC-T cell co-culture studies performed in collagen matrix, we used a similar approach we described previously (10, 46). Briefly, 270μL of bovine or rat-tail collagen solution at concentration of 1.7mg/mL was placed in a 48-well plate and solidified in the incubator for 45 minutes, then overlayed with the same volume of collagen solution embedded with T cells only, DC and T cells (at a ratio of 1:3), or DCs, T cells and 6ug of CPI Lys. After solidification in the incubator for 45 minutes, gels were overlayed with 300μL of Immunocult-XF Cell expansion media and further incubated at 37°C/5% CO_2_ for the indicated times. Cells were recovered with collagenase treatment as above for flow cytometry analysis.

### T cell proliferation assay

Day 5 HIV-infected CD4^+^ T cells were labelled with Tag-IT proliferation dye (Biolegend; cat # 425101) according to manufacturer’s instructions prior to the DC:T cell co-culture studies. Briefly, T cells were washed with PBS at 1500 rpm (400xg) for 10 mins and resuspended at 10^6^ cells/mL in a working dye solution of 5μM at 37°C/5% CO_2_ for 20 mins in the dark. The labelled CD4^+^ T cells were then washed with PBS and centrifuged at 1500 rpm for 10 mins. The pelleted cells were resuspended in Immunocult-XF Cell expansion media supplemented with rhIL-2 (5 μg/mL). To prevent additional rounds of HIV infection, 10 μM of the integrase inhibitor raltegravir, or a combination of 10 μM raltegravir (NIH HIV Reagents Program; cat # 11680), 20 μM emtricitabine (NIH HIV Reagents Program; cat # 10071) and 10 μM tenofovir (NIH HIV Reagents Program; cat # 10199) was added to culture throughout the experiment. To assess the proliferative capacity of infected T cells, Tag-IT labeled T cells were co-cultured with autologous DCs at a ratio of 1:5 (DC: T cell) in 24-well plate at different time points. In some studies, monocyte derived dendritic cells (MDDCs) were pulsed with 6μg/mL of protein pool consisting of Cytomegalovirus, Parainfluenza and Influenza (CPI, Immunospot; cat # CTL-CPI-001) and co-cultured with HIV infected T cells. At designated time points, proliferation of HIV latent T cells and HIV-exposed, double-negative T cells was assessed by flow cytometry. For neutralization studies, MDDCs were first pretreated with FcR blockade (Biolegend; cat #422302) to prevent FcR-mediated opsonization and incubated at 4°C for 20 mins. Afterwards, cells were washed, pelleted, and incubated with 10μg/mL of either isotype or neutralizing antibodies and incubated at 4°C for 30 mins. Antibody-neutralized DCs were co-cultured with HIV infected T cells in 24 well-plates, as described above. Blocking antibodies used: 10μg/mL each of anti-human CD80 (R&D; cat #MAB140), anti-human CD86 (R&D; cat #MAB141-100), anti-human HLA DR/DP/DQ (BD Bioscience; cat#555556) and 25 μg/mL of anti-human HLA-A/B/C (Biolegend; cat#311428). After 7 days of co-culture, culture supernatant was collected for further analysis, and cells were harvested for phenotypic analysis by flow cytometry. In some studies, SEB was added at a final concentration of 0.5µg/mL.

### TCR sequence analysis of expanded HIV-infected T cells

Tag-IT labelled, day 5 HIV Nef-2A-CRIMZs infected CD4^+^ T cells were co-cultured with autologous DCs in the presence or absence of CPI antigens at a ratio of 1: 5 (DC: T cell) in 24-well plate for 10 days in the presence of ART. At day 10 post co-culture, HIV latently infected CD4^+^ T cells were stained with CD3 and CD4 fluorochrome conjugated antibodies (Biolegend), fixed with 1.5% PFA and were sorted for proliferated and non-proliferated Zsgreen^+^ CD4^+^ T cells via flowcytometry (FACSAria III, BD Biosciences). Genomic DNA was isolated according to the manufacturer’s protocol from the sorted cells with the use of QIAamp DNA Formalin Fixed, Paraffin-Embedded Advanced UNG kits (QIAGEN, Maryland United States). Afterwards, 50ng of DNA was processed using the Illumina Ampliseq TCRB-SR Panel primers to amplify the CDR3 region, then libraries were indexed and purified using the Illumina Ampliseq library protocol. A pooled library was created by diluting samples to 2nM and then run on a MiSeq using a 300 cycle V2 kit at a final concentration of 7pM, with 10% PhiX v3 sequencing control. Raw sequences were processed through the command line to trim sequences and remove low quality sequences (<Q30 scores) using Cutadapt and trimmomatic packages, and aligned to the TCRB region using MiXCR (version 3.0.13). Immunarch R script was used to analyze data, and ggplot was used for visualization. T cell receptor antigen specificity was inferred by running the amino acid CDR3ß sequence through the IEDB (Immune Epitope Database and Tools) resource analysis lab – TCRMatch, using the trim sequence and a high (>.97) filtering cutoff for specificity.

### Near full length proviral sequencing analysis

CD4^+^ T cells isolated from 3 people without HIV were expanded and infected with HIV Nef-2A-CRIMZs and subsequently co-cultured with autologous DCs in the presence of CPI lysates, as described above. Proliferating (Tag-IT^+^) and non-proliferating (Tag-IT^neg^) DN and ZsGreen^+^CD4^hi^ T cells were sorted after 7 days of culture and processed for downstream PCR amplification. Briefly, T cells were resuspended in DirectPCR Lysis Reagent (Viagen), with 0.4mg/mL proteinase K and incubated at 55°C for 16h followed by 10 min at 95°C to inactivate proteinase K. Total HIV DNA was measured by PCR (112) in order to assess the number of HIV copies per uL of lysate. Near full-length amplification of HIV genomes were performed on lysates diluted to reach a concentration of 2 copies of HIV per well using a nested PCR as described previously (113). Briefly, HIV genomes were pre-amplified using Invitrogen Platinum SuperFi II MasterMix with 0.2mM of each primer (5’-CTCTGGTAACTAGAGATCCCTC-3’ and 5’-TCACACAACAGACGGGCACACACTACTT-3’) using a 25 cycle 3-step PCR protocol (initial denaturation at 98°C for 30s, denaturation at 98°C for 10s, annealing at 60°C for 10s, elongation at 72°C for 7 min, followed by a final extension at 72°C for 5 min). The pre-amplified products were diluted 1:9 with water, and 5uL were used in a second round of amplification, for 30 cycles (same cycling condition as the first amplification), using Invitrogen Platinum SuperFi II MasterMix with 0.2mM of each primer (5’-AGTCAGTGTGGAAAATCTCTAG-3’ and 5’-GCACTCAAGGCAAGCTTTATTGAGGCTTA-3’). The lengths of the amplicons were verified on a 0.8% agarose gel. Reactions selected for sequencing were purified with Ampure XP beads

(Beckman Coulter) according to manufacturer’s protocol, before proceeding to library preparation using Oxford Nanopore Technologies Native Barcoding Kit 96 V14. The sequencing was performed on R10.4.1 flow cells on a Gridion Sequencer (Oxford Nanopore Technologies). Analysis was performed using Nanopore wf-amplicon workflow. Integrity was assessed by MAFFT alignment against the HIV Nef-2A-CRIMZs reference sequence.

### Expansion, HIV infection and characterization of HIV latency in primary CD4^+^ T cell clones

SARS-CoV-2 specific CD4^+^ T cell clones (P155, P156) and autologous B cell lines isolated from patients were obtained from the Ostrowski lab and cultured as described (61). Both T cell clones were infected with HIV Nef-2A-CRIMZs at 0.1 MOI and assessed for infection by flow cytometry. 5 days post-infection, T cells were labelled with Tag-IT proliferation dye and co-cultured with autologous B cells in the presence of increasing cognate peptide concentration (P155; LRGHLRIAGHHLGRC and P156; LRIAGHHLGRCDIKD). All co-culture studies were performed using the indicated cognate peptide amount in a final volume of 1mL media (eg 50ng/mL). At day 2, the activation status of latent T cells (CD69, HLA-DR, PD-1) was assessed by flow cytometry, and proliferation (Tag-IT) and Gag p24 expression was measured at day 8 of co-culture. In some studies, anti-PD-1 blockade (3 μg/ml) prior to T:B co-culture in the presence of 100ng/mL peptides was performed.

## Statistical analysis

One-way ANOVA (Tukey’s multiple comparisons test and Sidak’s multiple comparisons test), Unpaired Student’s *t*-test, and Mann-Whitney U test (with non-normally distributed data) were used for data analysis using Prism 8 (GraphPad). The p-values from statistical analyses are shown in each graph. Differences were considered as not significant when p-values were greater than 0.05. Figures and illustrations were prepared using Adobeâ Illustrator and Biorenderâ.

## Data availability

Any additional information required to reanalyze the data reported in this paper is available from the lead contact upon request.

## Acknowledgements

We would like to thank Dr. Christine Zhang from the Flow core facility at the University of Manitoba. Antiretroviral drugs were purchased from the NIH HIV Reagent Program. This work was supported in part by the Canadian Institute for Health Research (CIHR) Project grant (PJT 155951 to T.T.M.), the CanCure Enterprise (HIG-133050 to T.T.M., N.C.) and Research Manitoba (T.T.M.).

## Notes

### Competing Interest Statement

The authors have declared no competing interest.

