## Supplementary Figures 1-9 for "Antigen stimulation drives clonal expansion of latent CD4^+^ T cells using a full-length HIV latency reporter"

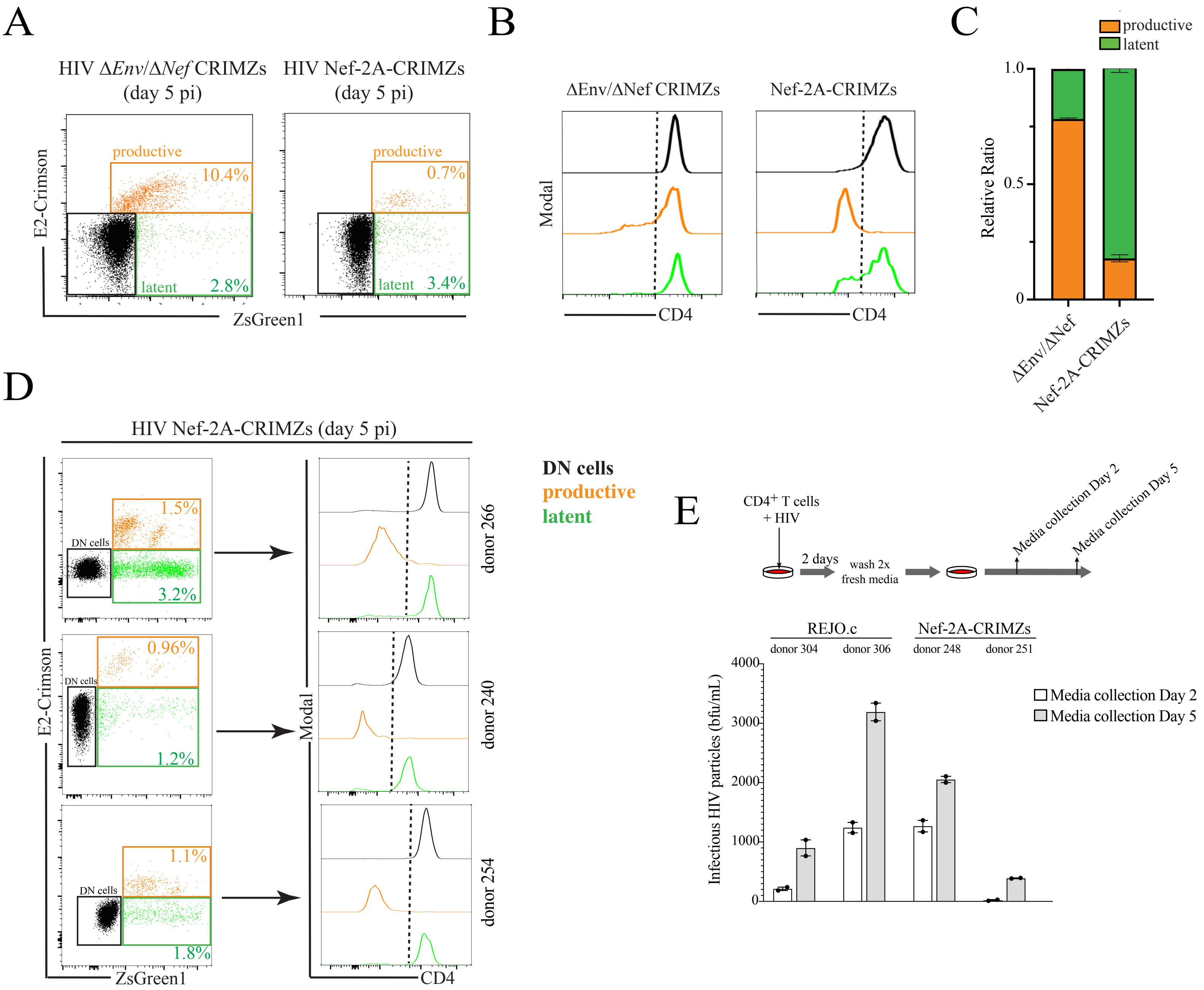


**Supplementary Figure 1: Functional Nef and Env expression in the HIV Nef-2A-CRIMZs reporter.** (A) Cultured Tcm were infected with the indicated dual-fluorescent HIV reporter and infection analyzed at day 5 by flow cytometry. (B) Comparative analysis of cell surface CD4 expression between the three T cell populations. Black line = uninfected, orange line = productive infection, green line = latent infection. (C) Relative ratio of productive and latently-infected T cells between the two reporter systems. Representative data from experiments performed in 3 healthy donors. (D) Distribution of DN, productive and latent T cell populations and their CD4 expression levels across three donors are shown. (E) T cells were infected with either T/F HIV REJO.c or Nef-2A-CRIMZs for 2 days, washed extensively to remove viral innoculum and culture supernatant collected at day 2 and day 5. Infectious HIV particles were measured using MAGI.CCR5 indicator cells and expressed as blue-forming units (bfu) per milliliter.

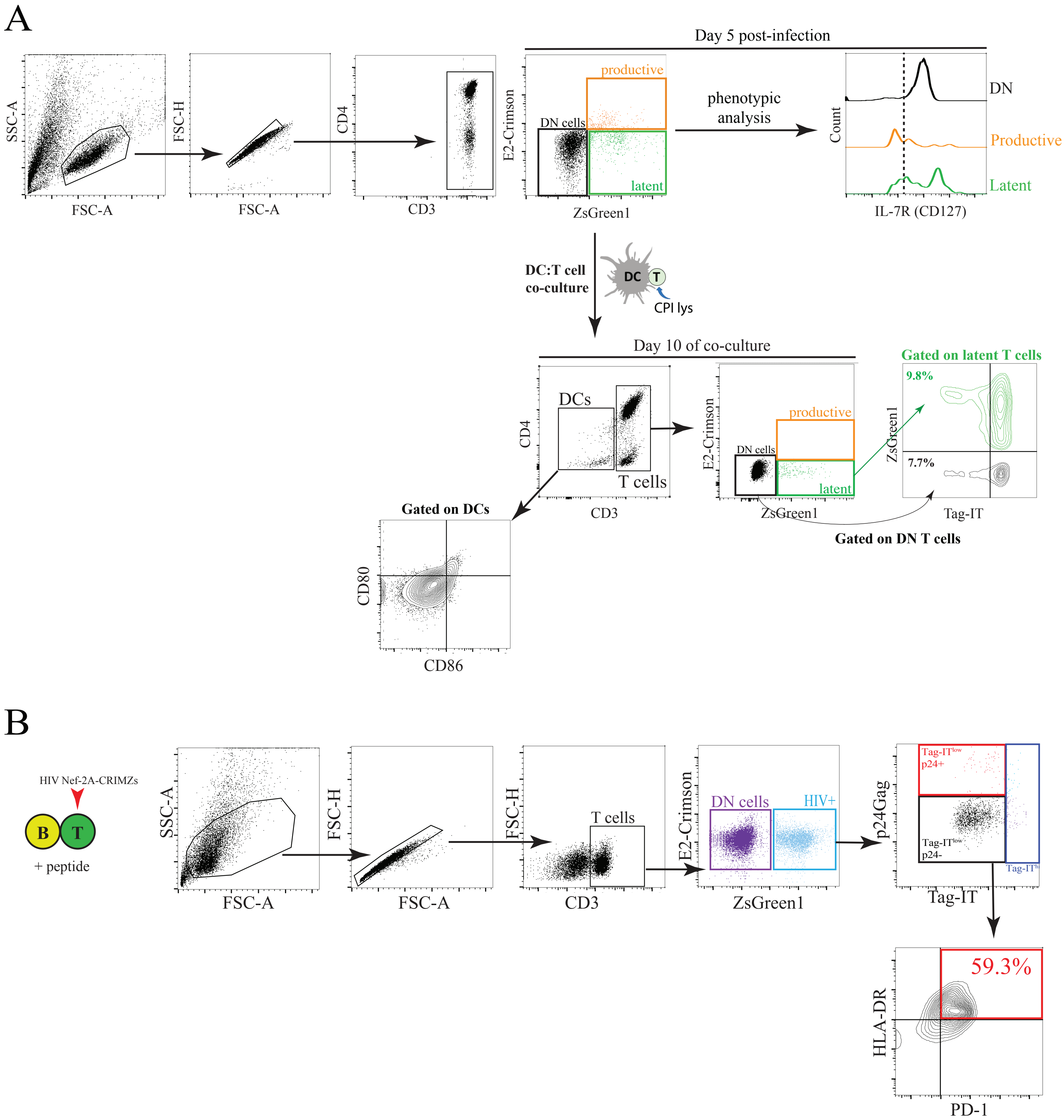


**Supplementary Figure 2: Flow cytometry gating strategy to identify HIV-infected T cell populations.** (A) Primary central memory CD4+ T cells were infected with HIV Nef-2A-CRIMZs and analyzed after 5 days by flow cytometry. Gating strategy to identify T cell populations (DN, productive, latent) for phenotypic analysis is shown. In some studies, infected T cells were co-cultured with autologous DCs in the presence of viral antigens: gating strategy in these co-culture studies is shown. Gating strategy on latent and HIV-exposed DN T cells is shown. (B) Gating strategy for T:B co-culture studies to identify proliferating, latent and DN T cells. Phenotypic analysis of both populations are shown below.

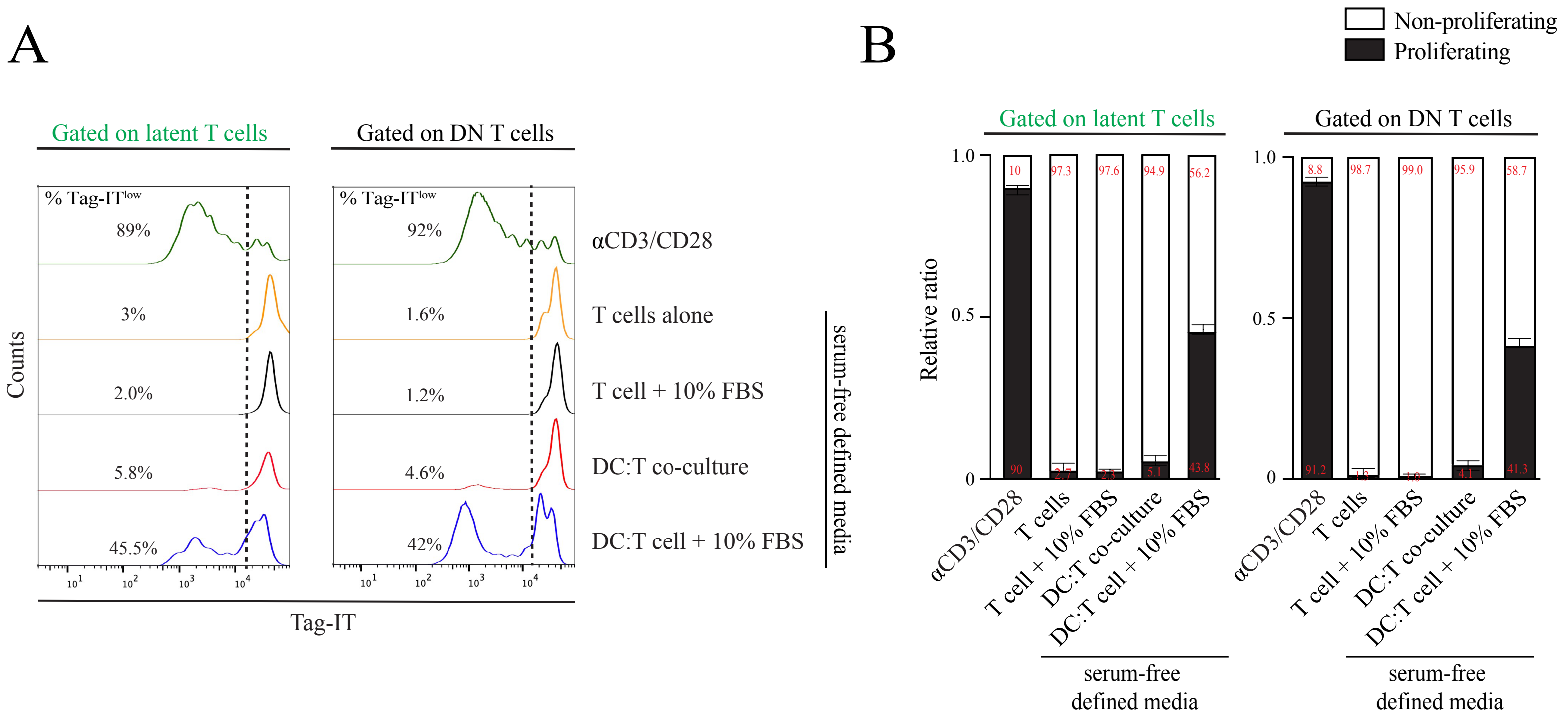


**Supplementary Figure 3: FBS in media induces latently-infected CD4+ T cell proliferation in a DC-dependent manner.** (A) Uninfected or latently-infected T cells were either left alone or co-cultured with DCs in the presence of FBS in serum-free defined media. Proliferative responses after 4 days was measured by flow cytometry. (B) Relative ratio of proliferating/non-proliferating T cells in response to the indicated culture conditions. Graphical representation of a representative donor out of 3 independent experiments. DN=double negative.

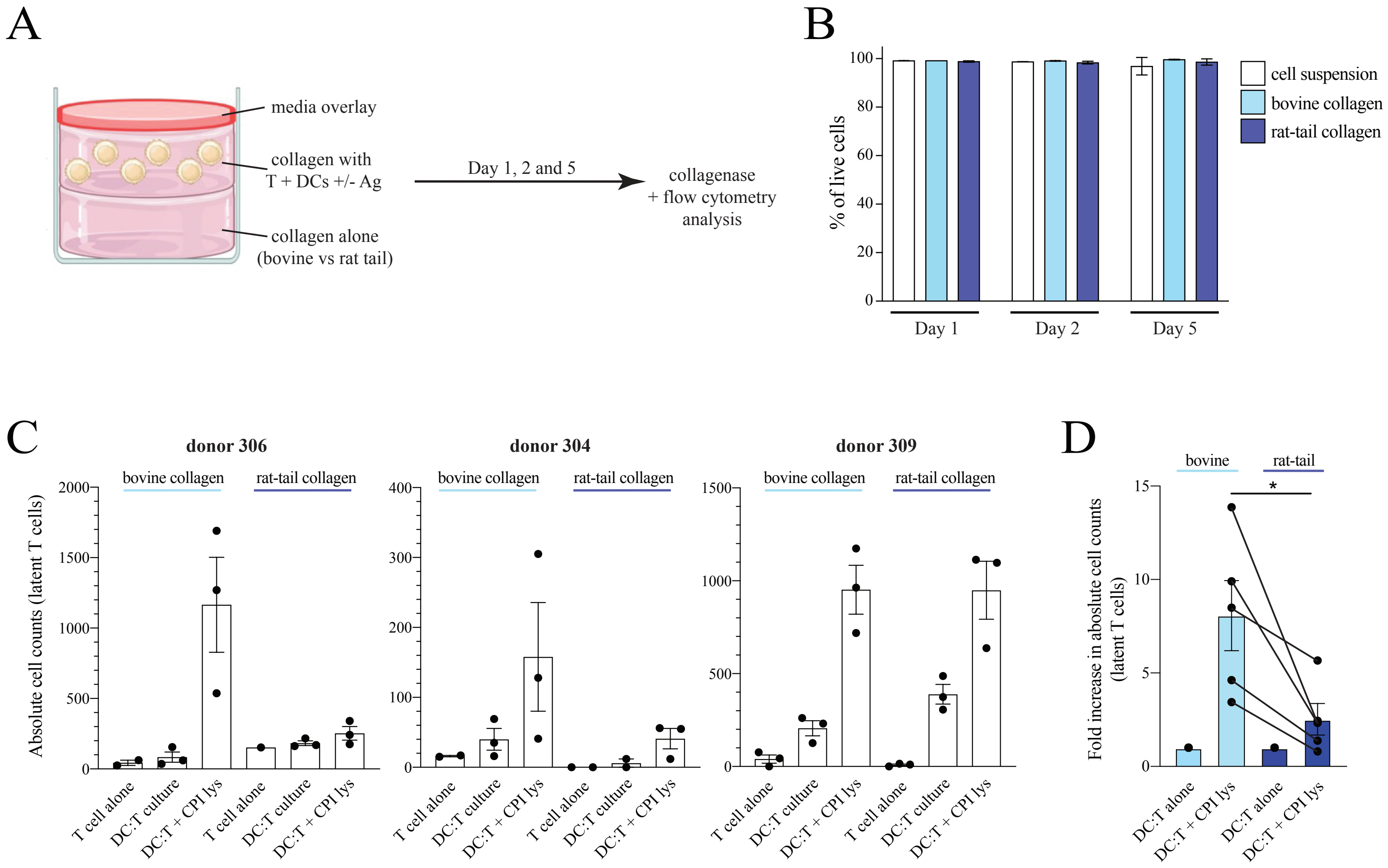


**Supplementary Figure 4: DC:T cell co-culture in 3D collagen.** (A) Autologous T cells and DCs were embedded into either bovine or rat-tail collagen at 1.7mg/mL and overlayed with media. A bottom layer of the corresponding collagen without cells was placed to create a physical buffer with the plastic surface. Cells were placed in the incubator for up to 5 days, treated with collagenase and analyzed by flow cytometry. (B) Total cell viability measured using a live/dead fixable dye. (C) Absolute cell counts of latent T cells across three representative donors shown. Conditions were T cell alone, DC:T cells alone and DC:T + CPI lys. Each dots represent biological replicates for each experiment. (D) Cumulative data from 5 donors, showing fold increase in cell counts in each culture condition. * $p<0.05$. Statistical analysis used = Mann Whitney nonparametric U test.

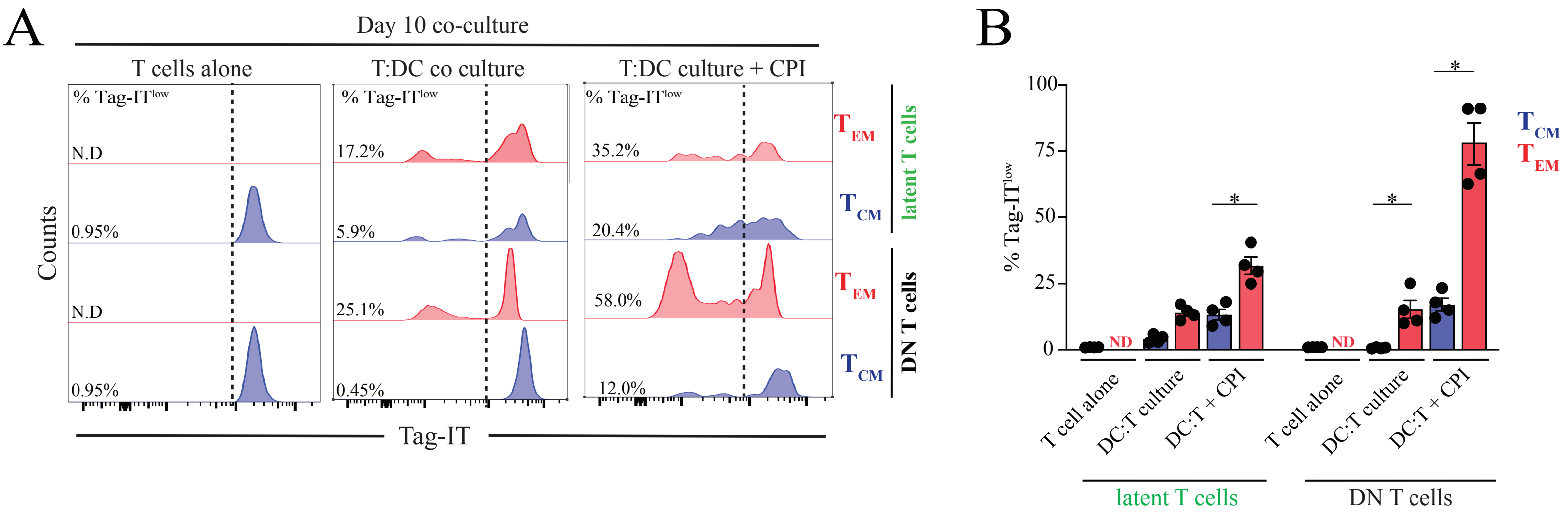


**Supplementary Figure 5: Phenotypic analysis of infected T cells in the presence or absence of autologous DCs.** (A) Tag-IT dilution of effector and central memory-like T cells in each culture condition. Percentages indicate Tag-IT-low population. (B) % Tag-IT-low values across subsets and conditions. ND=not detected. *p<0.05, Tukey's multiple comparisons test.

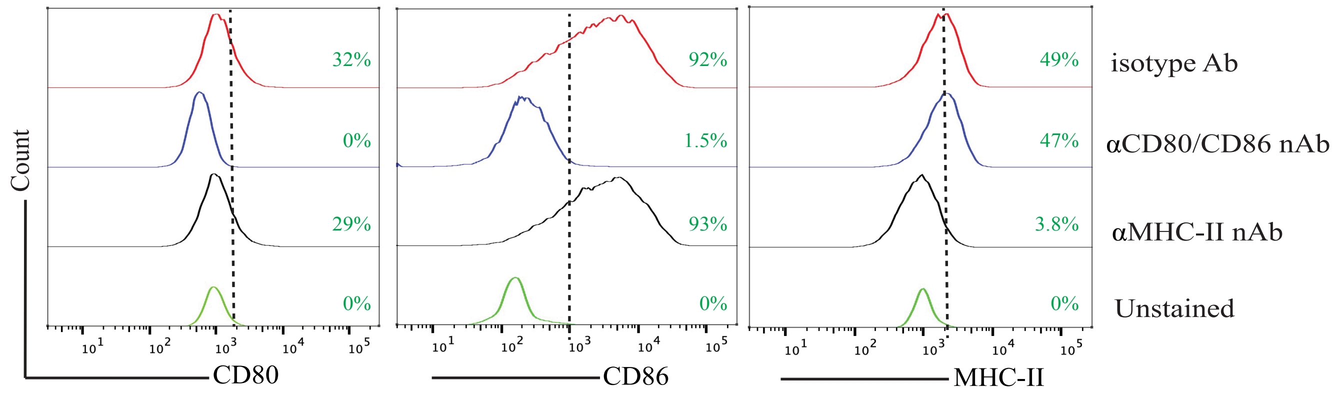


**Supplementary Figure 6: Validation of blocking antibodies used in Figure 4.** Cell surface CD80, CD86 and MHC-II expression after pre-treatment with the indicated blocking antibodies at day 8 is shown. Percentages indicate positive staining based on the dotted line.

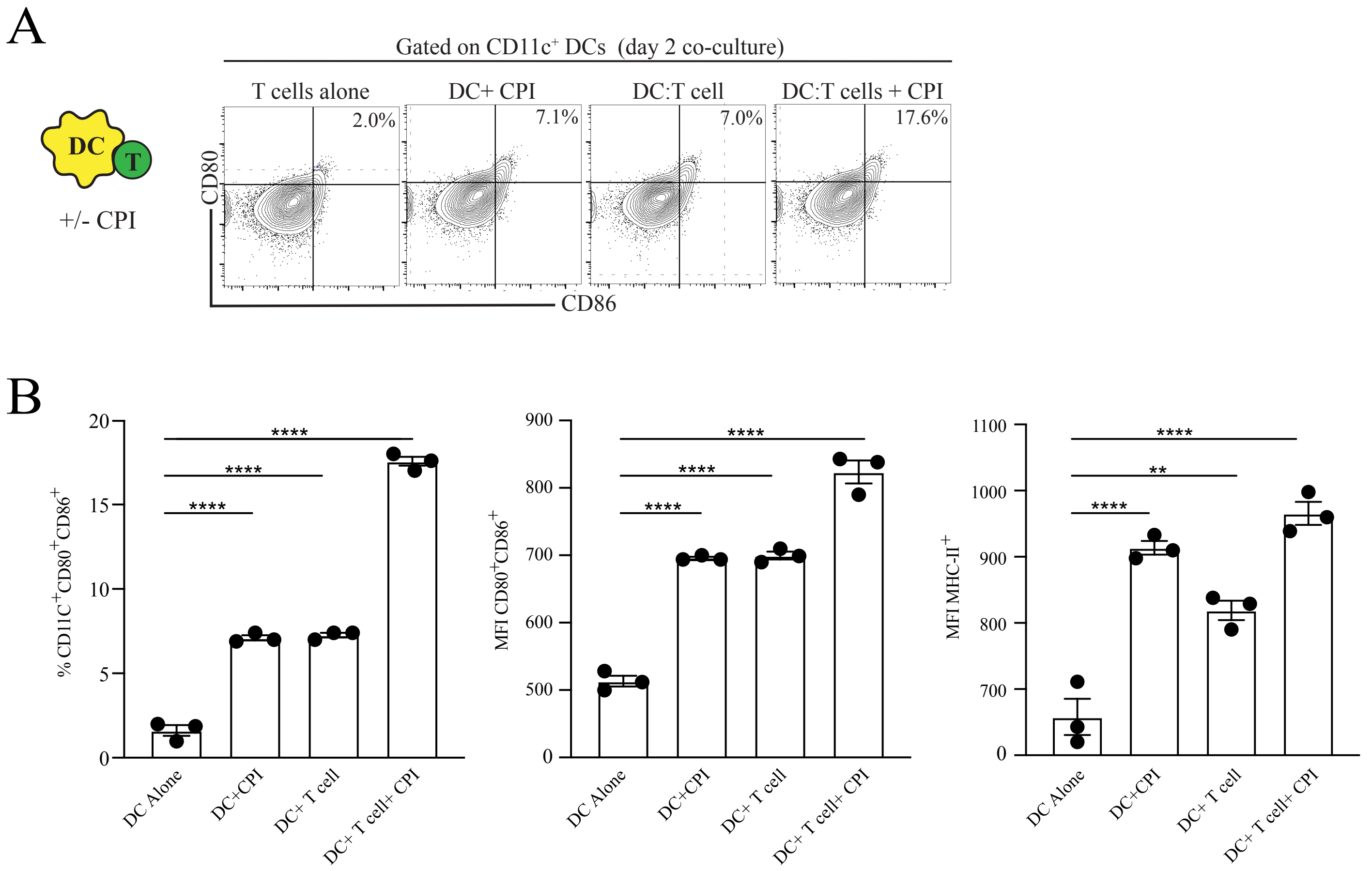


**Supplementary Figure 7: Phenotypic analysis of MDDCs during T:DC co-cultures.** (A) Autologous DC:T cell cultures in the presence or absence of antigen (CPI pool) were harvested and DCs stained for MHC-II, CD80 and CD86 expression by flow cytometry at day 2. (B) Phenotypic analysis of $CD11c^{+}$ DCs in the indicated culture conditions. Graphical representation of a representative donor out of 3 independent experiments is shown on the right. **$p<0.002$, ****$p<0.0001$. Statistical analysis used = Tukey's multiple comparisons test

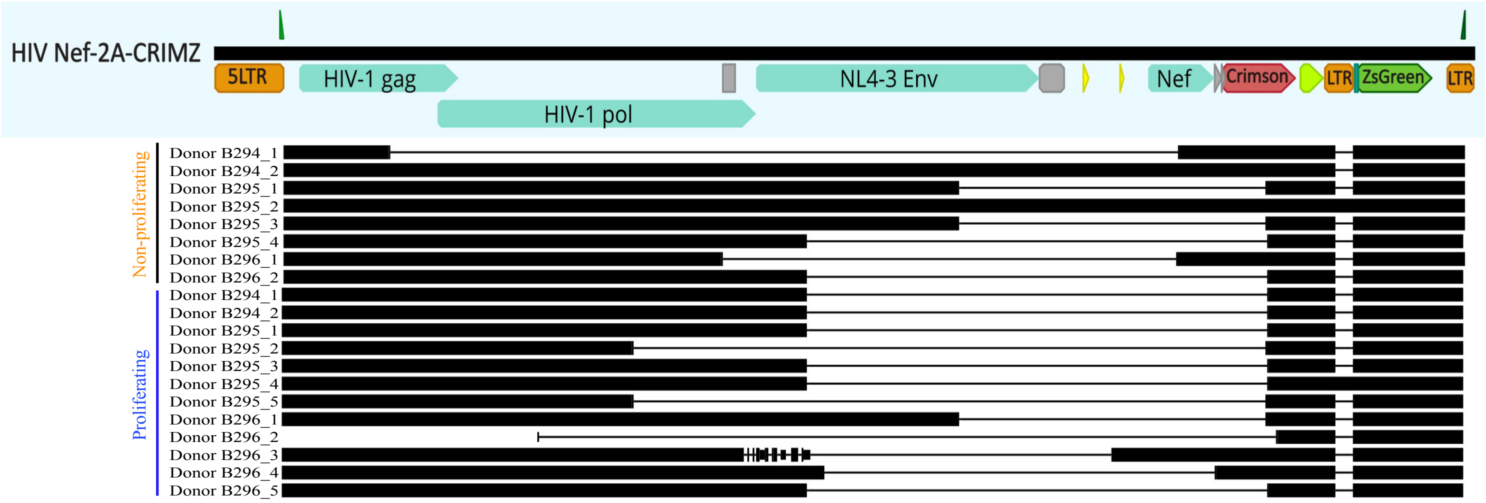


**Supplementary Figure 8: List of different defects found in 18 sequences from sorted proliferating and non-proliferating T cells.** Sequences are aligned on the HIV Nef-2A-CRIMZs genome.

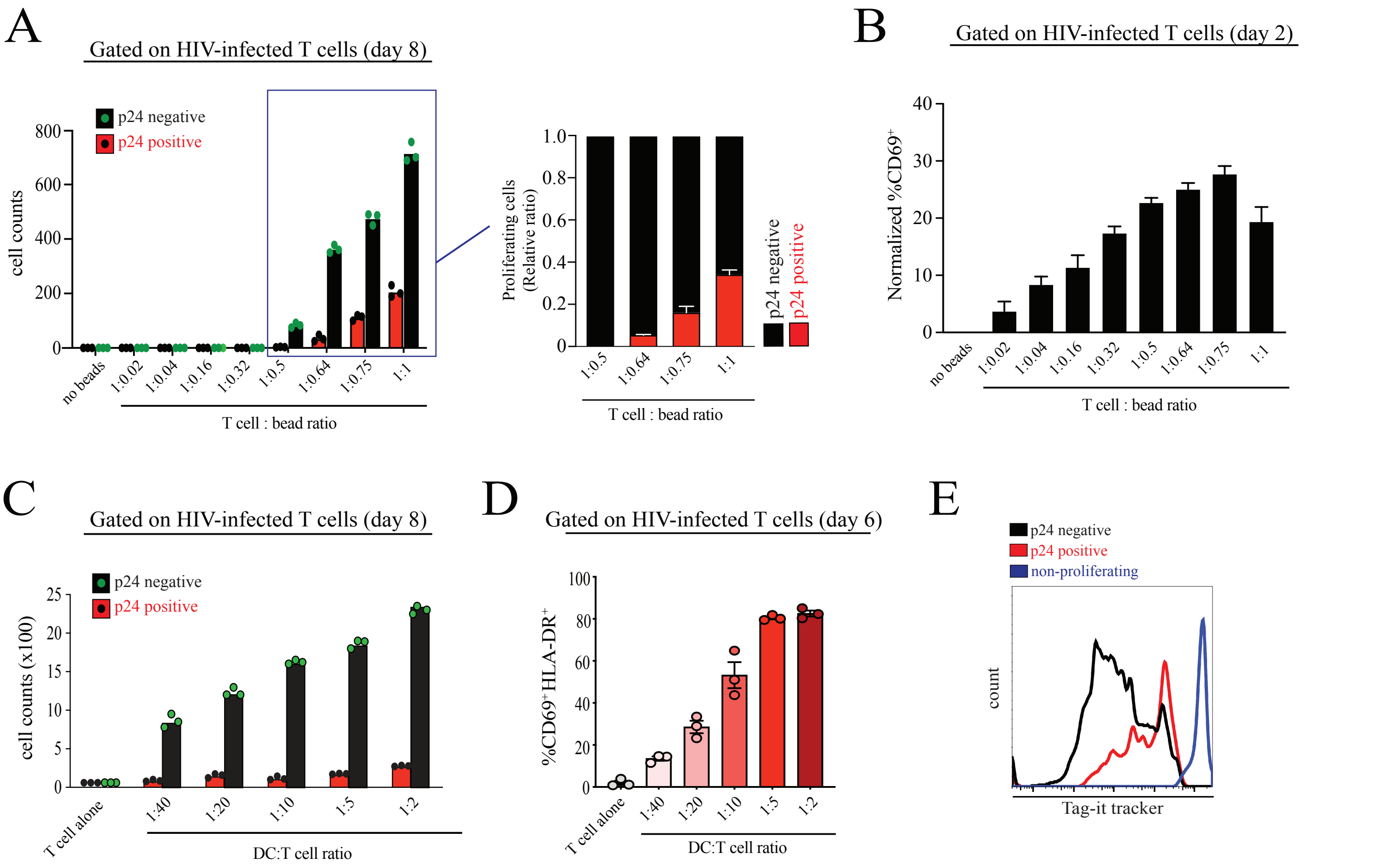


**Supplementary Figure 9: Interplay between TCR signaling strength and latent T cell activation, proliferation and proviral transcription.** (A) Absolute HIV-infected T cell counts after stimulation with the indicated anti-CD3/CD28 conjugated bead doses for 8 days. T cells were stained for Gag p24 and absolute cell counts enumerated by flow cytometry. Proliferative responses were measured using the Tag-it tracker dye. (B) Cell surface CD69 expression (day 2) is shown. Expression levels were normalized to no bead conditions. (C) Absolute HIV-infected T cell counts after co-culture in varying DC:T cell conditions. Varying DC:T cell co-culture conditions modulate proliferative responses of HIV infected T cells. (D) Cell surface CD69 and HLA-DR expression (day 6) is shown under different DC:T cell ratio. (E) Proliferative responses in p24 expressing or non-expressing T cells at the 1:10 DC:T cell ratio. Gag $p24^{+}$ T cells displayed reduced proliferative capacity over time.
